# The self-renewing lymphoid-biased hematopoietic stem cells support long-term lymphopoiesis in mice

**DOI:** 10.64898/2026.11.02.763559

**Authors:** Yao Ma, Xinying Li, Xialin Li, Lisha Wang, Yinglian Xu, Jinglei Zhai, Jinhong Wang, Yu Lan, Bo Zhou, Tao Cheng, Hideo Ema, Fang Dong

**Affiliations:** State Key Laboratory of Experimental Hematology, National Clinical Research Center for Blood Diseases, Haihe Laboratory of Cell Ecosystem, Institute of Hematology & Blood Diseases Hospital, Chinese Academy of Medical Sciences & Peking Union Medical College, Tianjin 300020, China; Tianjin Institutes of Health Science, Tianjin 301600, China; Department of Stem Cell & Regenerative Medicine, Peking Union Medical College, Tianjin, China; Key Laboratory of Multi-Cell Systems, Shanghai Institute of Biochemistry and Cell Biology, Center for Excellence in Molecular Cell Science, Chinese Academy of Sciences; University of Chinese Academy of Sciences, Shanghai 200031, China

**Keywords:** Lymphoid-biased HSC (Ly-biased HSC), EPCR^+^MPP6 (EMPP6), EPCR^+^MPP5 (EMPP5), single-cell assays, paired daughter cells (PDCs)

## Abstract

Lymphopoiesis is profoundly impaired during ageing largely due to ageing-susceptible changes of hematopoietic stem cells (HSCs). How to maximize the preservation of lymphopoiesis during ageing remains unknown and whether there exists a specific population with life-long lymphoid potential not intensively affected by ageing is of substantial scientific and translational significance. Here, we identified a self-renewable, lymphoid-biased hematopoietic stem cells (Ly-biased HSCs) subset within the EPCR+CD150^−^CD48^−^LSK (HSC2) population, phenotypically termed EPCR^+^MPP6 (CD34^−^CD135^−^HSC2), that not only displayed comparable multi-lineage potential with HSC in young mice, but also maintained similar lymphoid-biased reconstitution during aging. The lymphoid-biased characteristics of EPCR^+^MPP6 were also supported by immune associated program at transcriptome and proteomic levels. Notably, EPCR^+^MPP6 and HSCs rarely generated one another following the first cell division in vitro and under homeostasis in vivo indicating the most likely independent relationship between them and an earlier-than-thought bifurcation of the lymphoid differentiation pathway, which may function as a backup source to support lymphopoiesis. Collectively, this newly defined Ly-biased HSCs represent a promising cellular target for both mechanistic studies and therapeutic strategies concerning age-related immune decline.

**Highlights:**

1. The self-renewable lymphoid-biased hematopoietic stem cells (Ly-biased HSCs) are identified in mice.
2. Unlike HSCs, the Ly-biased HSCs maintain substantial lymphoid potential during ageing.
3. Ly-biased HSCs initiate an alternative lymphoid differentiation pathway which is likely independent from conventional multipotent HSCs.

## INTRODUCTION

The deterioration of lymphopoiesis and therefore adaptive immunity is one of the characteristics of ageing, leading to increased vulnerability to infection in the elderly.^1–4^ Although variant explanations have been proposed, most of them are boiled down to the ageing-related inherent changes in hematopoietic stem cells (HSCs).^3–6^ Interestingly, alongside the ageing-susceptible HSC, emerging evidence has independently revealed existence of ageing-resilient populations with unaffected or only mildly compromised lymphoid output, which, for example, were non-expanded^6^ or exhibited “younger” functional and molecular features,^7^ or were with high mitochondrial mass.^8^ These findings raised the possibility of an alternative source of lymphopoiesis distinct from the one initiated by those ageing-susceptible HSCs.

Despite the controversy over step-wise or continuous models,^9^ hematopoietic hierarchy in general is composed of three branches, where the lymphoid lineage is distinctively separated from the myeloid and megakaryocyte/erythroid (MkE) lineages.^10^ Along with the great improvement in immunophenotypic purification, especially the use of two SLAM family markers CD150 and CD48,^11^ this bifurcation point has moved from common lymphoid progenitors (CLP) to lymphoid-primed multipotent progenitors (LMPP), and to the commonly acknowledged MPP4 at present.^12–21^ EPCR, a marker for long-term HSCs (LT-HSCs) in both embryonic and adult stages and an indicator of quiescence,^22–26^ was recently exploited to identify MPP Ly-I, which exhibited superior function than MPP4 and was transcriptionally more approximate to HSCs,^5^ raising the possibility of an even earlier bifurcation event. However, none of these reported progenitors are able to independently support long-term lymphopoiesis due to their lack of self-renewal potential.

We previously defined EPCR^+^CD150^+^CD48^−^LSK as HSC1 (functionally, LT-HSC), and parallelly found EPCR^+^CD150^−^CD48^−^LSK (HSC2) enriched for lymphoid ancestors with repopulating capacity.^27,28^ However, it remains unknown whether a more primitive lymphoid ancestor exists considering the great heterogeneity within HSC2. Here, we first confirmed that HSC2 enriched for the more primitive lymphoid-biased progenitors. Next, we found that HSC2 was composed of three subsets: EMPP6 (CD34^−^CD135^−^HSC2), EMPP5 (CD34^+^CD135^−^HSC2) and the previously reported MPP Ly-I (CD34^+^CD135^+^HSC2), which formed a functional and developmental hierarchy. Notably, unlike the aberrantly expanded aged HSC1 with compromised lymphoid readout during ageing, EMPP6 from aged mice only mildly expanded and displayed almost comparable lymphoid potential with their young counterparts. Moreover, the strong possibility of independent relationship between HSC1 and EMPP6 evidenced by pair daughter cell assay in vitro and lineage tracing in vivo indicated an earlier bifurcation of the lymphoid-lineage differentiation pathway, which may function as a backup source to support lymphopoiesis.

## RESULTS

### EPCR enriches for a more primitive lymphoid-biased population in CD150^−^CD48^−^LSK cells

EPCR^+^CD150^+^CD48^−^LSK cells (HSC1) was reported to enrich for the LT-HSC population.^26,28–31^ Since CD150^−^CD48^−^LSK cells can also be fractionated by EPCR (**Fig. 1A**), we hypothesized that these EPCR^+^ cells (HSC2) can similarly enrich for a more primitive lymphoid-biased population. To verify this, transplantation of HSC1, HSC2 and EPCR^−^CD150^−^CD48^−^LSK (DN) from GFP transgenic mice (CD45.2) was performed and injected to lethally irradiated recipients (total 9.0Gy, CD45.2). Only GFP^high^ cells were considered true donor-derived cells to exclude potential false positive cells sticked by GFP^+^ platelet (**Fig. S1A**).

**Fig. 1.**
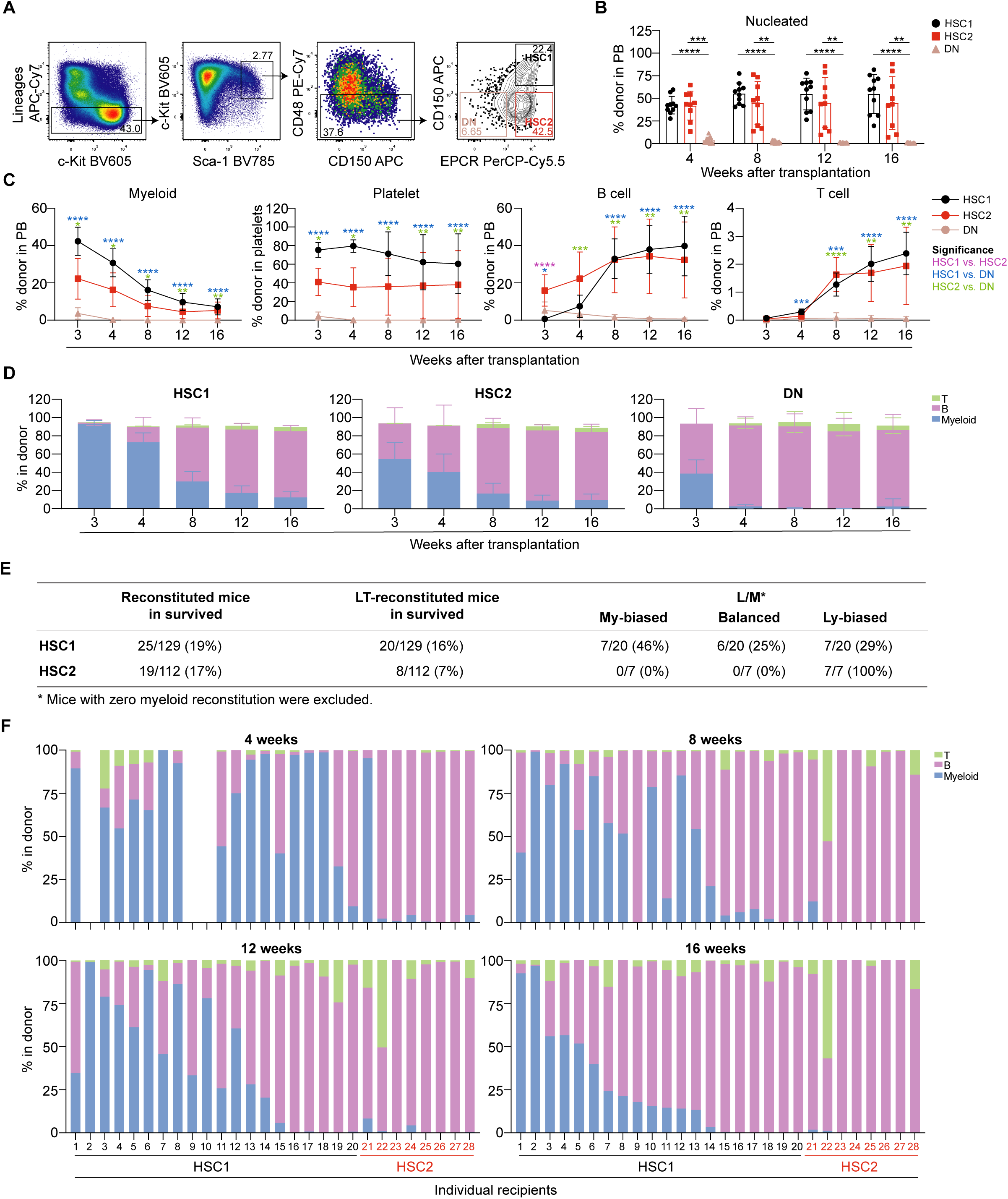
EPCR^+^CD150^−^CD48^−^LSK (HSC2) cells enrich for self-renewable cells compared to EPCR^−^ ones. **(A)** Flow cytometry profile of EPCR expression in CD48^−^Lineage^−^Sca-1^+^c-Kit^+^ (LSK) cells. **(B-C)** Donor-derived total nucleated **(B)**, myeloid, B cell, T cell reconstitution in PB and donor proportion in PB platelets **(C)** after transplantation. Data were collected from two independent experiments, HSC1, n = 10; HSC2, n = 9; DN, n = 12. **(D)** Myeloid, B cell and T cell reconstitution from same experiments with **(C)** but presented by proportion in donors. **(E)** Table showing detailed results of single-cell transplantation. **(F)** Myeloid, B cell and T cell reconstitution at 4, 8, 12 and 16 weeks presented by proportion in HSC1 and HSC2 donors after single cell transplantation. Data were shown as mean ± SD. \**P* ≤ 0.05, \*\**P* ≤ 0.01, \*\*\**P* ≤ 0.001, and \*\*\*\**P* ≤ 0.0001 (one-way ANOVA for Fig. 1B **and C**). See also Fig. S1.

As expected, DN barely reconstituted after transplantation, whereas HSC2 maintained a comparable level of reconstitution to HSC1 throughout 16 weeks (**Fig. 1B**). For lineage reconstitution (**Fig. S1B-C**), HSC2 exhibited a classic lymphoid-biased reconstitution where an early but transient myeloid/platelet (MyeP) wave was followed by stable B and T cell reconstitution **(Fig. 1C, D)**. Consistent results were obtained from the bone marrow (BM) and spleen (SP) of recipients 16 weeks after transplantation **(Fig. S1D, E).**

To address the heterogeneity within HSC2 cells, single-cell colony assay and single-cell transplantation were performed with HSC1 in parallel. HSC2 cells formed significantly fewer colonies than HSC1 (**Fig. S1F, G**). For colony types, neutrophils and/or macrophages (nm) colonies dominated in HSC2 cells, whereas colonies including neutrophils and/or macrophages, erythrocytes, and megakaryocytes (nmE/Mk) dominated in HSC1 (**Fig. S1H**). For single-cell transplantation assay, 20 out of 129 mice (16%) were long-term (16 weeks) reconstituted by HSC1 cells, while 8 out of 114 mice (7%) were long-term reconstituted by HSC2 cells (**Fig. 1E**). HSC2 cells exhibited poor myeloid and T cell reconstitution, but relatively better B cell reconstitution through 16 weeks (**Fig. 1F, S1I**). The lymphoid versus myeloid (L/M) ratio was further calculated to quantitively identify the lineage biases of donor cell ^32^ (**Fig. S1J**). Notably, while more than 70% of HSC1 clones were myeloid-biased (My-biased) and balanced, all HSC2 clones were classified as lymphoid-biased (Ly-biased) (**Fig. 1E**). Collectively, these results confirmed that EPCR could efficiently enrich for a more primitive, yet heterogeneous, lymphoid-biased population in CD150^−^CD48^−^LSK cells.

### HSC2 cells are a heterogeneous population composed of three immunophenotypically distinct subsets

To understand the molecular characteristics of HSC2, scRNA-seq was performed. HSC2 cells were clearly separated from HSC1, suggesting a distinct transcriptomic profile (**Fig. 2A**). Differentially expressed genes (DEGs) upregulated in HSC2 were largely related to the lymphoid lineage, such as *Cd52*, *Flt3*, *Tespa1*, *Cd27* and *Il17ra*. In contrast, downregulated genes in HSC2 cells included those associated with LT-HSC, such as *Pdzik1ip1*, *Sult1a1*, and *Mllt3*, as well as those related to myeloid and megakaryocyte lineages, such as *Cavin2*, *Esam*, *Gata2*, and *Selp* (**Fig. 2B**). Gene ontology (GO) terms of upregulated genes in HSC2 cells were predominantly associated with lymphocyte activation and differentiation, whereas those of downregulated genes were largely related to platelet functions (**Fig. 2C**). Notably, Gene Set Enrichment Analysis (GESA) of HSC2 based on previously reported signatures^33–36^ revealed similarity with short-term HSC (ST-HSC), consistent with previous observations that Ly-HSC overlapped with ST-HSC.^37^ Besides, HSC2 also shared similar features with MPP5^36^ and MPP4 (**Fig. 2D**).

**Fig. 2.**
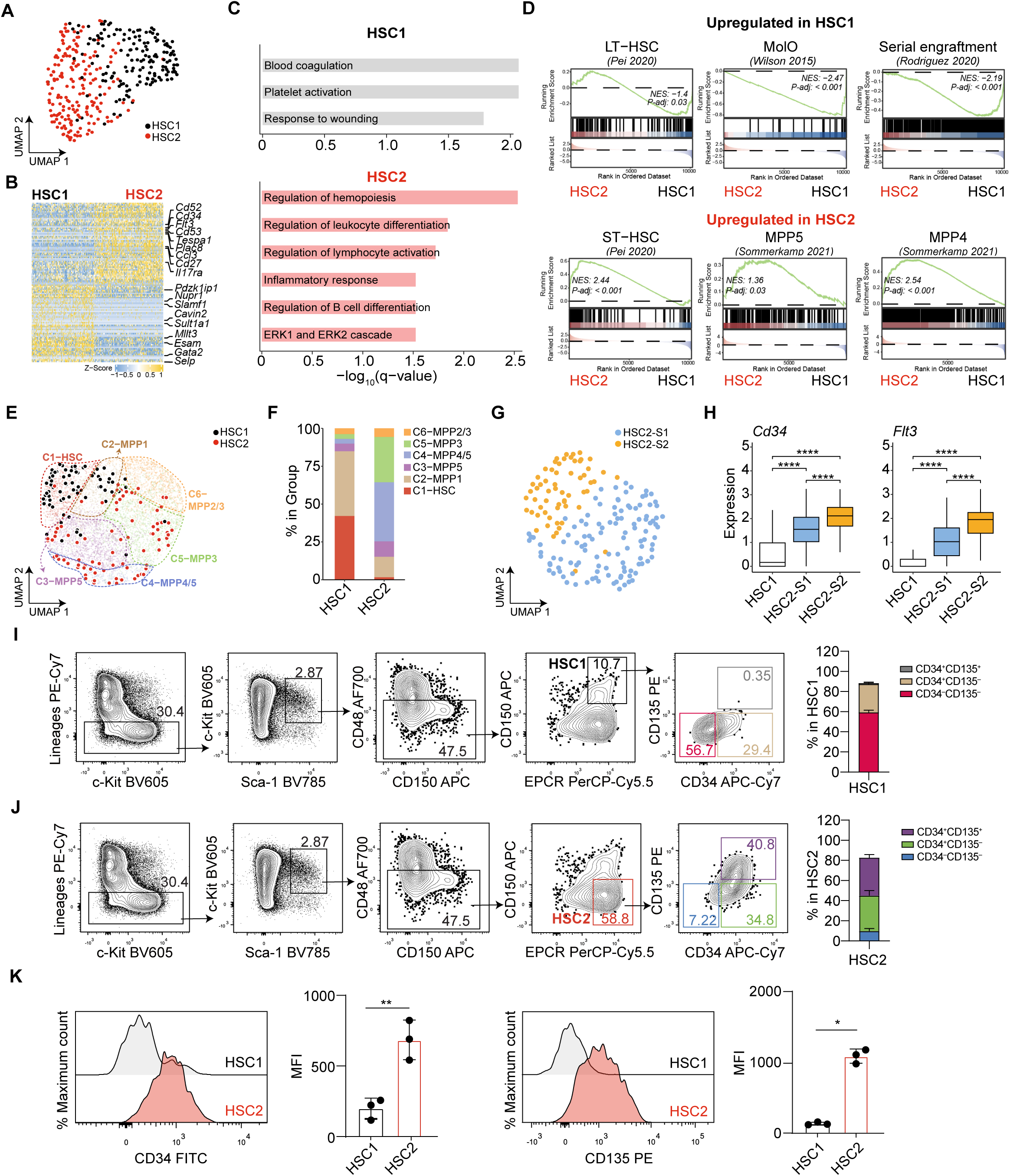
HSC2 displays lymphoid lineage signatures with heterogeneity. **(A)** Uniform Manifold Approximation and Projection (UMAP) visualization of 159 HSC1 and 177 HSC2 cells, collected from at least 3 mice. **(B)** Heatmap showing differentially expressed genes (DEGs) between HSC1 and HSC2. **(C)** Gene Ontology (GO) analysis of DEGs in HSC1 (above) and HSC2 (bottom). **(D)** Gene Set Enrichment Analysis (GSEA) using HSC-and MPP-related gene sets previously published.^33–36^ **(E)** UMAP plot highlighting specific distribution of HSC1 and HSC2 cells on the integrated data (LSK data previously published^36^ together with HSC1 and HSC2 data from this study). **(F)** Summary of cluster distribution in HSC1 and HSC2 cells based on the integrated data in Fig. 2E. **(G)** Subclusters further identified in HSC2 (HSC2-S1 and -S2). **(H)** Box plots showing expression level of *Cd34* and *Flt3* in HSC2-S1 and -S2 with HSC1 as control. **(I)** Representative flow cytometry profiles showing the CD34 and CD135 expression in HSC1 (left) and bar plot showing relative proportions (right). **(J)** Similar to **(I)** except in HSC2 cells. **(K)** Median fluorescence intensity (MFI) of CD34 and CD135 in HSC1 and HSC2 cells. Data were shown as mean ± SD. \**P* ≤ 0.05, \*\**P* ≤ 0.01, \*\*\**P* ≤ 0.001, and \*\*\*\**P* ≤ 0.0001 (wilcoxon test for Fig. 2B, H, unpaired Student’s t-test for Fig. 2K). See also Fig. S2.

To better delineate the heterogeneity of HSC2 cells, we integrated scRNA-seq dataset published previously^36^ with HSC1 and HSC2. Six clusters were identified, each assigned an identity of immunophenotypic HSCs or MPPs (**Fig S2A, B**). HSC2 displayed a more scattered distribution pattern than HSC1 (**Fig. 2E**). Specifically, HSC1 cells were predominantly distributed in C1-HSC and C2-MPP1, while HSC2 cells were mainly in C4-MPP4/5 and C5-MPP3 (**Fig. 2F**).

Two sub-clusters were further identified in HSC2, sub-cluster 1 (HSC2-S1) and sub-cluster 2 (HSC2-S2) (**Fig. 2G**). The most predominant features separating HSC2-S1 and -S2 were *Cd34* and *Flt3* (**Fig. 2H**), prompting us to investigate the corresponding flow cytometry profiles. Most HSC1 cells appeared to be in the CD34^−/low^CD135^−^ fraction (**Fig. 2I, S2C**). In contrast, HSC2 comprised CD34^−^CD135^−^, CD34^+^CD135^−^, and CD34^+^CD135^+^ cells (**Fig. 2J**). CD34^−^CD135^−^ and CD34^+^CD135^−^ cells overlapped with MPP6 and MPP5, respectively, despite that MPP6 and MPP5 contained both EPCR-negative and positive cells (**Fig. S2D**). CD34^+^CD135^+^ cells within HSC2 were almost identical to the recently reported lymphoid-committed MPP (MPP Ly-I).^5^ The median fluorescence intensity (MFI) of CD34 and CD135 of HSC2 was greater than that of HSC1 (**Fig. 2K**). In addition, comparison of EPCR expression levels revealed a gradual decrease from HSC1 to MPP4, except for MPP6 (**Fig. S2E**).

Together, these results suggest that HSC2, while displaying the lymphoid-lineage signatures, was a heterogeneous population that immunophenotypically overlapped with MPP6, MPP5 and MPP Ly-I.

### The three subsets in HSC2 form a functional ranking from the aspect of their self-renewal and multi-linage potential

Based on CD34 and CD135 expression, we further subdivided the HSC2 cells into three subsets: EPCR^+^MPP6 (EMPP6), EPCR^+^MPP5 (EMPP5), and MPP Ly-I (**Fig. S2D, 3A**). Cell cycle analysis was first performed using fluorescent ubiquitination-based cell cycle indicator (Fucci) mice with Cdt1-monomeric Kusabira-Orange 2 (mKO2) and monomeric geminin-Azami Green (mAG).^26,38,39^ mKO2^high^ cells are in the quiescent (G0) state, whereas mKO2^low^, mKO2^neg^, and mAG^+^ cells are in the G0/1, G1, S/G2/M states respectively. Although all three subsets exhibited a substantial degree of quiescence, EMPP6 was the most quiescent comparable with HSC1 **(Fig. 3B)**.

**Fig. 3.**
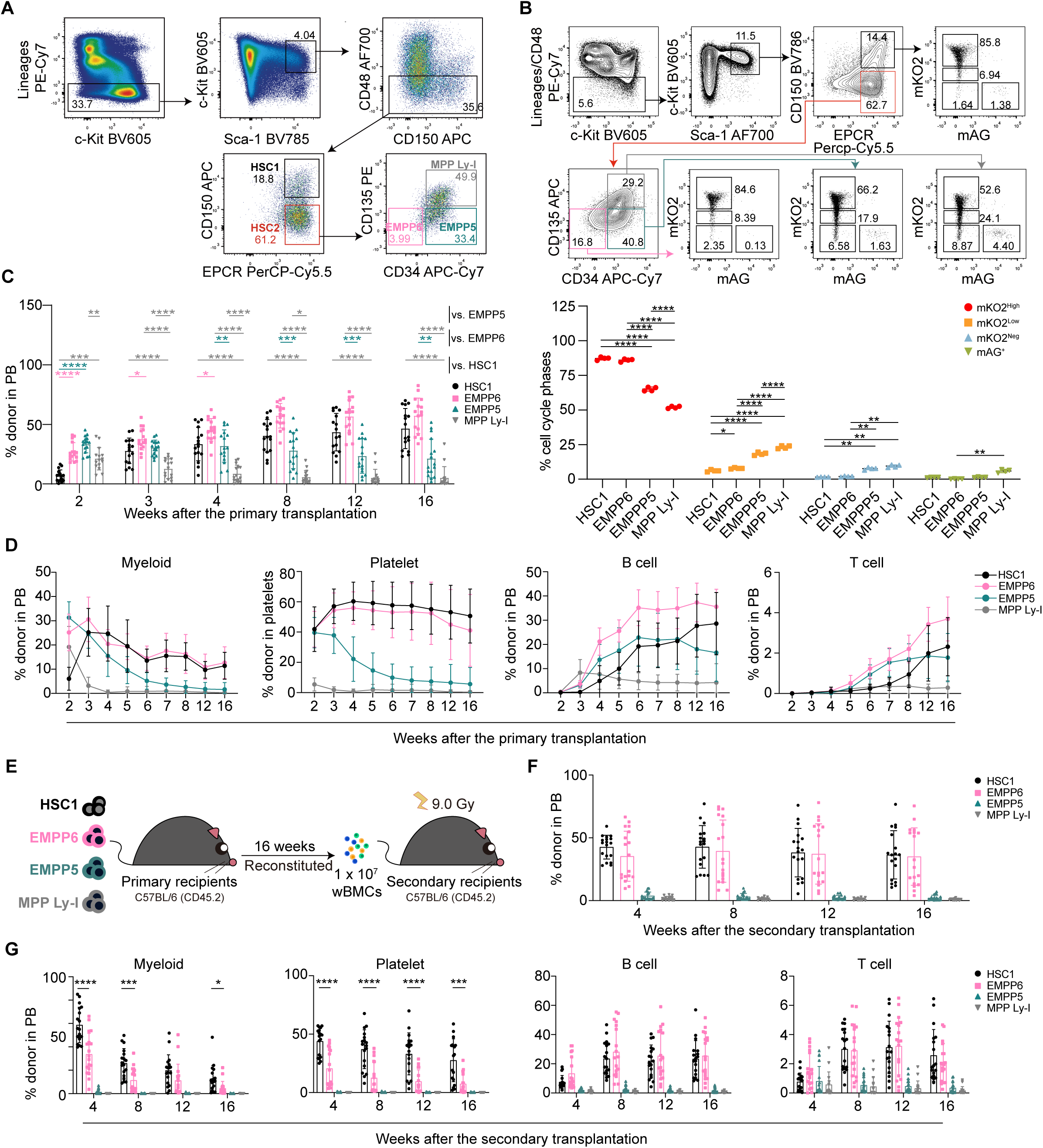
HSC2 subsets form a functional hierarchy. **(A)** Representative flow cytometry profiles of HSC2 subpopulations fractionated by CD34 and CD135 expression. **(B)** Representative flow cytometry profiles (above) and statistical analysis (bottom) of HSC1, EMPP6, EMPP5 and MPP Ly-1 cells in the BM using fluorescent ubiquitination-based cell cycle indicator (Fucci) mice. Data were shown from one out of two independent experiments, n = 4 for each group. **(C)** Donor-derived total reconstitution in PB after the primary transplantation. Data were collected from three independent experiments, HSC1, n = 16; EMPP6, n = 16; EMPP5, n = 16; and MPP Ly-I, n = 15. **(D)** Donor-derived myeloid cell, B cell, T cell reconstitution in PB and donor proportion in PB platelets after the primary transplantation. **(E)** Schematic plot showing workflow of the secondary transplantation. **(F)** Donor-derived reconstitution in PB after the secondary transplantation. Data were collected from three independent experiments, HSC1, n = 18; EMPP6, n = 18; EMPP5, n = 18; and MPP Ly-I, n = 16. **(G)** Donor-derived myeloid cell, B cell, T cell reconstitution in PB and donor proportion in PB platelets after the secondary transplantation. Data were shown as mean ± SD. \**P* ≤ 0.05, \*\**P* ≤ 0.01, \*\*\**P* ≤ 0.001, and \*\*\*\**P* ≤ 0.0001 (One-way ANOVA for Fig. 3B, C; only HSC1 and EMPP6 were compared using unpaired Student’s t-test in Fig. 3F, G). See also Fig. S3.

Given that CD34 and CD135 are well established markers for HSCs and MPPs, we assumed a functional hierarchy among the three subsets and performed competitive transplantation. The total reconstitution level in peripheral blood (PB) progressively decreased along EMPP6, EMPP5 and MPP Ly-I, but was comparable between EMPP6 and HSC1 (**Fig. 3C**). As for mature lineages, EMPP6 displayed significantly faster myeloid (2 weeks) and B cell (3 weeks) reconstitution than HSC1, and still greater B cell reconstitution during 4 to 8 weeks, followed by similar levels of reconstitution in all lineages until 16 weeks. EMPP5 and MPP Ly-I, however, showed faster but transient myeloid reconstitution, and MPP Ly-I barely reconstituted platelets. Both EMPP5 and MPP Ly-I were able to maintain the lymphoid-lineage reconstitution, but to a lesser extent (**Fig. 3D, S3A**). Consistent reconstitution patterns were observed in BM and SP at 16 weeks (**Fig. S3B, C**). During the secondary transplantation, comparable level of total reconstitution was still revealed between EMPP6 and HSC1, whereas EMPP5 and MPP Ly-I barely reconstituted, suggesting that EMPP6 was the only subset able to self-renew (**Fig. 3E, F**). However, EMPP6 displayed significantly reduced MyeP reconstitution compared to HSC1 (**Fig. 3G**).

We also examined donor-derived HSC and MPP populations in BM 16 weeks after both the primary and the secondary transplantation. To distinguish between HSC1 and MPP1, we used CD34^−^CD135^−^CD150^+^CD48^−^LSK instead of HSC1 as traditionally defined HSC (tradHSC) (**Fig. S3D-F, Supplemental Table S1**). After the primary transplantation, although EMPP6 could generate both tradHSC and MPP1, their proportions were significantly less than those derived from HSC1 (**Fig. S3G)**. After the secondary transplantation, however, donor-derived tradHSC, MPP1, EMPP5, and MyeP-lineage progenitors (i.e., =CMP, GMP, and MkP) were significantly reduced from EMPP6 than HSC1, while no statistical differences were detected for donor-derived EMPP6 and lymphoid-lineage progenitors (i.e., MPP Ly-I, MPP4 and CLP) between two groups (**Fig. S3H**).

Collectively, these results suggest that EMPP6, EMPP5 and MPP Ly-I contributed to the lymphoid lineages in a functional ranking, where EMPP6 lay at the top, representing the most primitive lymphoid-lineage ancestor.

### EMPP6 is functionally defined as lymphoid biased HSCs with self-renewal ability at the clonal level

To further confirm the functional identity of EMPP6, single-cell serial transplantation was performed on HSC1 and EMPP6 (**Fig. 4A**). The LT-reconstitution efficiency of EMPP6 (18%) was slightly higher than that of HSC1 (12%). Eleven out of 12 (92%) EMPP6 clones with detectable myeloid reconstitution at 16 weeks were classified as Ly-HSC in contrast to 4 out of 14 (29%) HSC1 clones (**Fig. 4B**). Notably, the only EMPP6 clone classified as Bal-HSC showed an L/M ratio of 9.9, extremely close to the Ly-HSC cutoff. Most EMPP6 clones showed only transient MyeP reconstitution (< 12 weeks). Therefore, the long-term reconstitution of EMPP6 clones was mainly contributed by considerably maintained B and T cells (**Fig. 4C-E**).

**Fig. 4.**
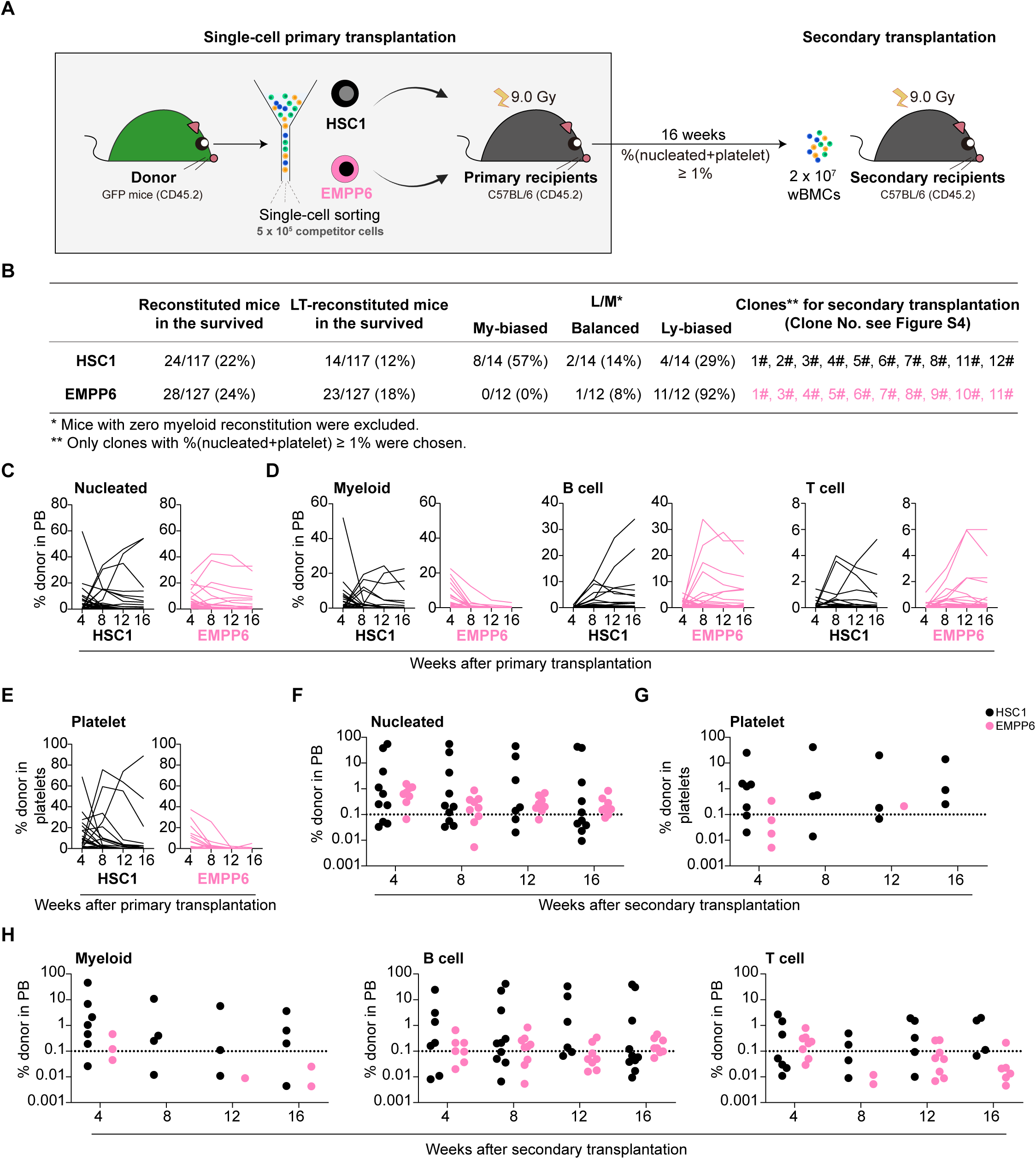
EMPP6 displays self-renewal ability at the clonal level. **(A)** Schematic plot illustrating the experimental design of single-cell primary and secondary transplantation. **(B)** Table showing the number of reconstituted mice and long-term (LT) reconstituted mice in the number of survived mice, L/M ratio-based clone classification and clones for secondary transplantation. **(C)** Donor-derived nucleated reconstitution in PB. All reconstituted recipients were shown. Each line represents one recipient. HSC1, n = 24; EMPP6, n = 28. **(D)** Donor-derived myeloid, B cell, and T cell reconstitution in PB. **(E)** Donor proportion in PB platelets. **(F)** Donor-derived nucleated reconstitution in PB following the secondary transplantation. Recipients with reconstitution > 0.001% were shown and the dash line indicated the threshold for reconstitution (> 0.1%). **(G)** Donor proportion in PB platelets following the secondary transplantation. **(H)** Donor-derived myeloid, B cell, and T cell reconstitution in PB following secondary transplantation. See also Fig. S4.

To decide the lineage potential for each clone, we defined four types of clones based on the reconstituted lineages (**Fig. S4A**). Four weeks after transplantation, EMPP6 contained all four types of clones, while HSC1 lacked “B/T” clones. Sixteen weeks after transplantation, while “B/T” clones dominated in EMPP6, a small proportion (10.7%) remained to be “PBMT” clones (**Fig. S4B**), indicating very rare maintained multi-lineage potential of EMPP6 after the primary transplantation, in addition to major clones that either quickly lost the MyeP potential or restricted to the lymphoid-lineages throughout life.

To investigate the self-renewal activity of the most durable clones in EMPP6, the secondary transplantation was performed (**Fig. 4A**). While some EMPP6 clones maintained the lymphoid reconstitution till 16 weeks following the secondary transplantation, none of them displayed discernable MyeP reconstitution (**Fig. 4F-H**). These results suggested that EMPP6 was mostly consisted of functional Ly-biased HSC clones with self-renewal potential but lymphoid-biased reconstitution compared to HSC1.

### EMPP6 displays immune-enriched features and a lower aging score compared with HSC1

To investigate the transcriptomic differences among EMPP6, EMPP5 and MPP Ly-I, scRNA-seq was performed **(Fig. S5A)**. Very few DEGs were discovered between either two groups (**Supplemental Table S2**). GSEA results showed that EMPP6 significantly upregulated inflammation/immune-related terms (TNFα signaling via NFκB, Interferon gamma/alpha response) but downregulated proliferation-related terms (E2F targets, G2M checkpoint, Myc targets) (**Fig. S5B**), consistent with cell cycle analysis (**Fig. 3B**). The three subsets also displayed a gradually increased scoring of lineage signatures, with EMPP5 more prominent in the megakaryocyte/erythroid lineages than MPP Ly-I (**Fig. S5C**). These results suggested a more primitive transcriptomic profile of EMPP6, a multipotent but more differentiated profile of EMPP5, and a more lymphoid lineage-committed profile of MPP Ly-I.

To learn the differentiation preferences of and the possible hierarchy relationship among HSC1, EMPP6, EMPP5 and MPP Ly-I, trajectory inferring was performed (**Fig. S5D**). While both HSC1 and EMPP6 could differentiate towards the myeloid lineage, EMPP6 exhibited clearer differentiation preferences towards the lymphoid lineages and HSC1 towards MkE. Notably, some EMPP6 clones were mixed with MPP Ly-I and CLP, suggesting there exist lymphoid-restricted progenitors in EMPP6, consistent with results of single-cell transplantation. The apex was mostly taken up by HSC1 and EMPP6 cells, followed gradually by EMPP5 and MPP Ly-I, suggesting a hierarchical order along HSC1/EMPP6, EMPP5 and MPP Ly-I. Surprisingly, most EMPP6 cells showed clear separation from HSC1 afterwards, not like EMPP5, MPP1 or MPP2 which were partially or completely downstream of HSC1, indicating an alternative lymphoid differentiation pathway initiated by most EMPP6 clones, if not all, alongside HSC1.

Therefore, we examined the molecular characteristics distinguishing EMPP6 from HSC1. Genes upregulated in EMPP6 were mainly related to: 1) Cytoskeleton, e.g., *Myl10*, *Cfl1*, *Arpc1b*, thymosin family (*Tmsb10*, *Tmsb4x*, *Ptma*); 2) Stress and activation, e.g., *Plac8*, *Ier2*, *Ramp1*; 3) Immune-related, e.g., *Ltb*, *Ifitm1, Ifitm2*. Genes downregulated in EMPP6 were mainly related to: 1) Reported LT-HSC markers, e.g., *Sult1a1*, *Npm1*, *Mmrn1*, *Tgm2*, *Apoe*, *Trim47*, *Clec1a*; 2) Megakaryocyte lineage, e.g., *Itga2b* (encoding CD41), *Itgb3* (encoding CD61); 3) Metabolism, e.g., *Serinc3*, *Sord*, *Acadl*, *Upp1*, *Aldh1a1* (**Fig. 5A and Supplemental Table S2**). Consistently, GSEA results from both transcriptomic and proteomic data suggested similar features of EMPP6 with an upregulation of immune/inflammation/cytoskeleton related pathways and a downregulation of metabolism related pathways (**Fig. 5B, C and Supplemental Table S3**).

**Fig. 5.**
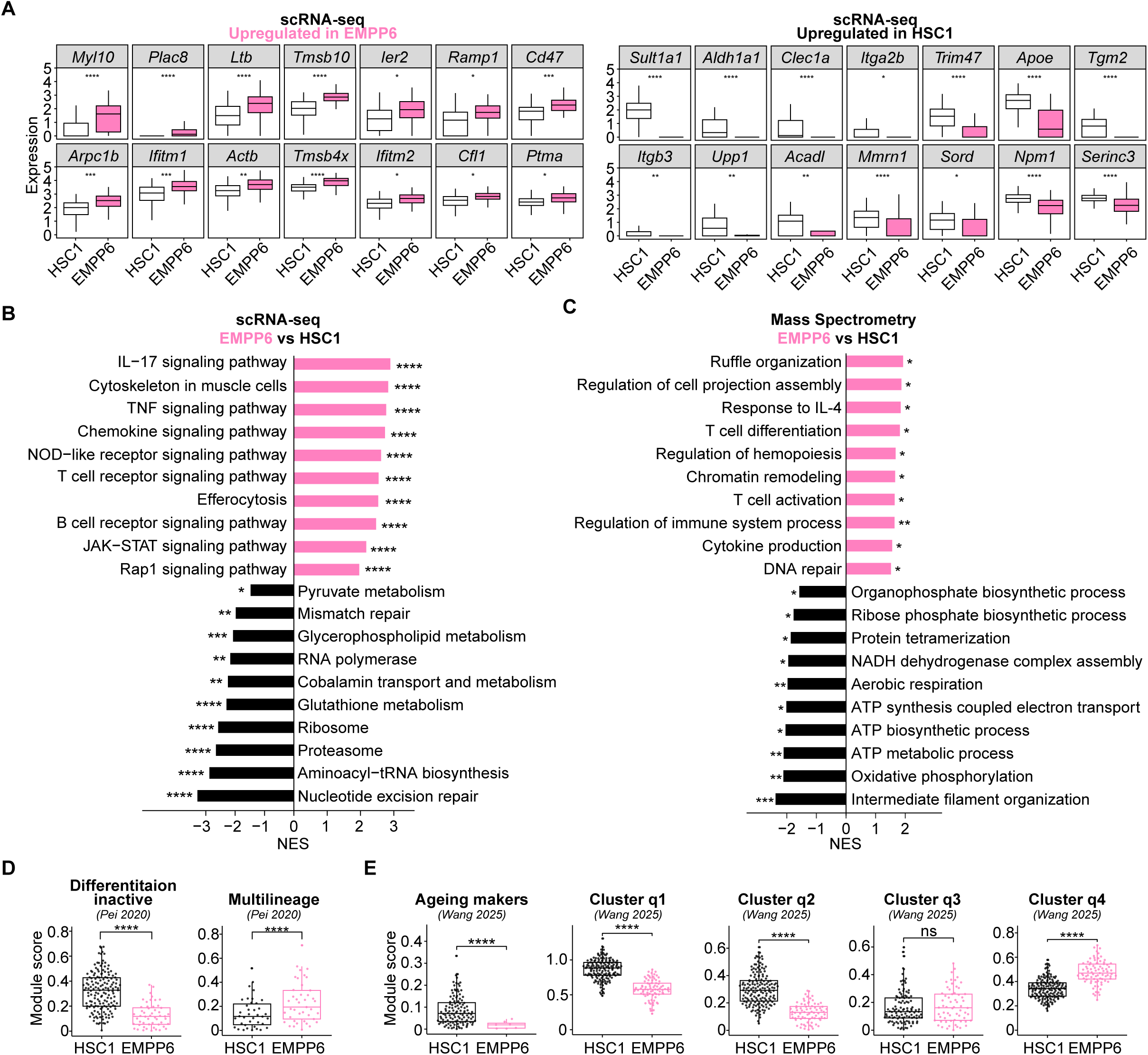
EMPP6 enriched for immune-related features while displaying a lower ageing score. **(A)** Differentially expressed genes between HSC1 and EMPP6 cells. Data were shown as mean ± SD. \**P* ≤ 0.05, \*\**P* ≤ 0.01, \*\*\**P* ≤ 0.001, and \*\*\*\**P* ≤ 0.0001 (wilcoxon test). **(B)** GESA on scRNA-seq data using the KEGG database between HSC1 and EMPP6 cells. **(C)** GESA on mass spectrometry data using the GO-BP database between HSC1 and EMPP6 cells. **(D)** Enrichment scoring using HSC-related gene sets.^33–35^ **(E)** Enrichment scoring using ageing-related gene sets.^7,56^ In the original article, cluster q1, q2, and q3 were identified from aged HSCs, whereas q4 was from young HSCs. See also Fig. S5 and Supplemental Table S2 to S3.

To investigate the role of EMPP6 more specifically in the hematopoietic settings, we calculated enrichment scores of EMPP6 and HSC1 with published gene sets.^33–35^ Not surprisingly, compared to HSC1 that exhibited magnificent LT-HSC and Low-output signatures, EMPP6 mainly enriched for ST-HSC and High-output clones (**Fig. S5E, F**). Consistent with this, EMPP6 also scored significantly higher in the “multilineage” gene set (**Fig. 5D**), which were suggested to have stronger correlation with ST-HSC,^35^ again confirming the multipotent nature of EMPP6. Given the negative expression of CD150 and obvious lymphoid-lineage bias of EMPP6, we next examined its association with ageing. Interestingly, EMPP6 exhibited a significantly lower ageing score than HSC1 (**Fig. 5E**), corroborating the correlation between CD150 expression and ageing.^7^

Collectively, these data demonstrated that EMPP6 represented a highly quiescent and multipotent population with immune-enriched features both in transcriptomic and proteomic level. Notably, despite both populations being derived from young mice, EMPP6 exhibited a lower aging score than HSC1, suggesting that EMPP6 might be better equipped to cope with age-associated decline.

### EMPP6 maintains robust lymphoid potential during ageing

The quiescent state and lymphoid-biased fate of EMPP6 prompted us to investigate how EMPP6 functionally changed with ageing. Although both the proportion and the absolute number of EMPP6 significantly increased in aged mice, they did not change as drastically as HSC1 (10.5-fold increase in HSC1 versus 5-fold in EMPP6 in **Fig. 6A**, 17.8-fold increase in HSC1 versus 5.3-fold in EMPP6 in **Fig. 6B**). Transplantation of these cells (**Fig. 6C**) revealed magnificently impaired reconstituting ability of aged HSC1 (A-HSC1) compared to young counterpart in PB and 16-week SP (though not in BM), in contrast to maintained or mildly reduced reconstitution level of aged EMPP6 (A-EMPP6) compared to young counterpart, but still much higher than A-HSC1 (**Fig. 6D, S6A and B**). Specifically, although the myeloid outputs were maintained by all groups in PB, BM and SP (**Fig. 6E, S6A and B**), the lymphoid output (B and T cells) from A-HSC1 was dramatically decreased compared to that of young group, which supported the myeloid-biased hematopoiesis of HSC1 with aging. In contrast, A-EMPP6 performed significantly better in lymphoid-lineage reconstitution than A-HSC1, including Pro B, follicular B and marginal-zone B cells and central memory T cells analyzed at 16 weeks in BM and SP, indicating its stability in keeping the inherent lineage bias which was only mildly affected by ageing-related stress (**Fig. 6E-G, S6C-F**). BM HSPC analyses after primary transplantation revealed reduced contribution of A-HSC1 to tradHSC, EMPP5, MPP Ly-I and MPP1-4 than Y-HSC1 (**Fig. S6G**). However, no statistical significance was found between Y-EMPP6 and A-EMPP6 except for CLP (**Fig. S6G**). PB results from the secondary transplantation showed comparable reconstitution of A-EMPP6 to Y-EMPP6 (**Fig. 6H, I**), indicating similarly self-renewal capacity of EMPP6 maintained from young to aged adulthood. Notably, A-EMPP6 still exhibited comparable B precursor reconstitution with Y-EMPP6 and Y-HSC1 (**Fig. 6J**), indicating sustained ongoing lymphopoiesis rather than the cumulative existence of long-lived lymphocytes.

**Fig. 6.**
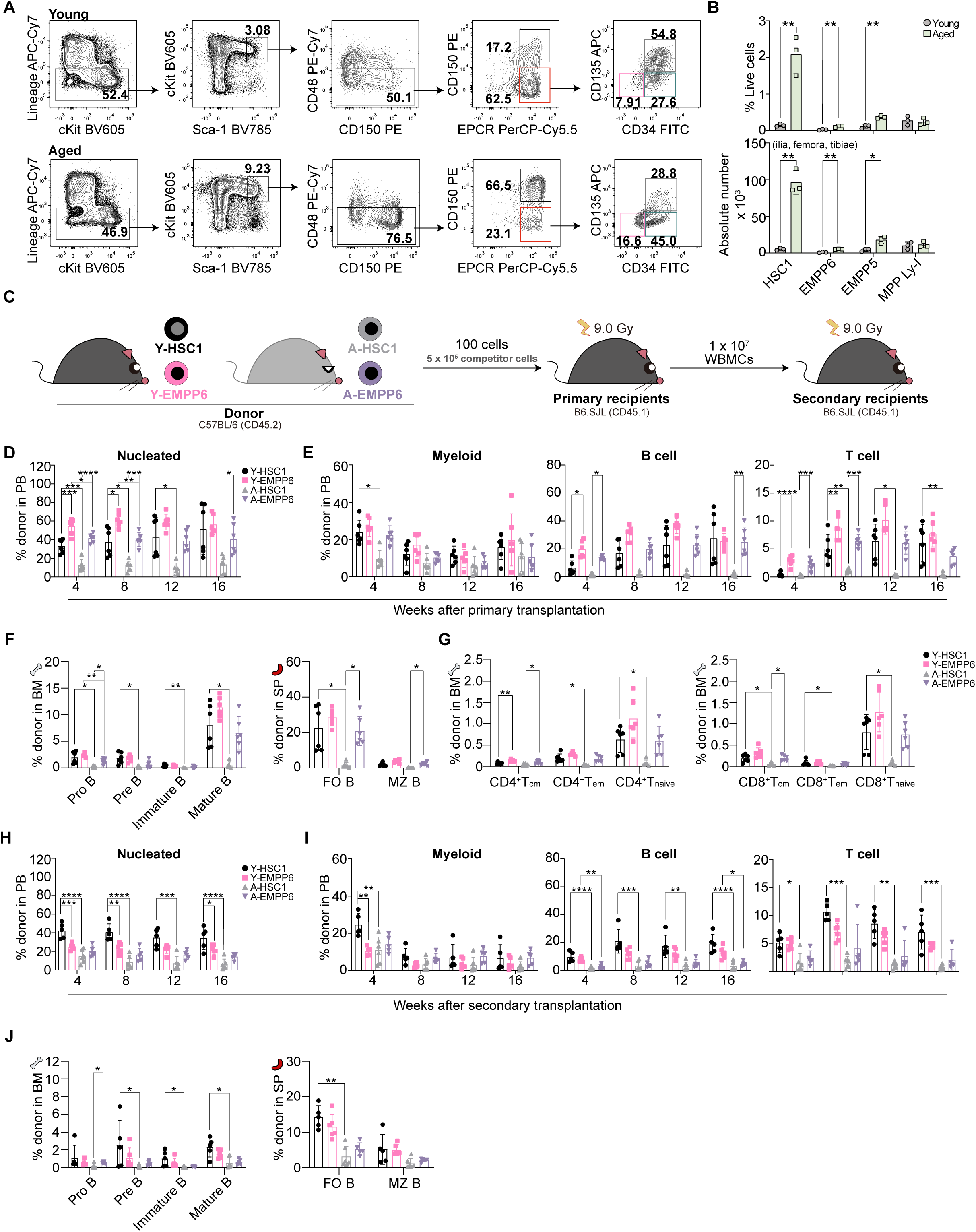
EMPP6 was of higher functional fitness than HSC1 in dealing with age-related stress. **(A)** Representative flow cytometry profiles of the indicated populations in young and aged mice. **(B)** Proportion (above) and the absolute number (bottom) of indicated populations in young and aged mice. Data were shown from one representative experiment, n = 3 in both groups. **(C)** Experimental design for the primary and secondary transplantation. **(D)** Donor-derived nucleated reconstitution in PB after the primary transplantation. Data were shown from one representative experiment, n = 6 in both groups. **(E)** Donor-derived myeloid, B and T cell reconstitution in PB. **(F)** Reconstitution of donor-derived B-cell subsets in 16-week BM (left) and SP (right) of recipients. **(G)** Reconstitution of donor-derived CD4^+^T (left) and CD8^+^T (right) subsets in 16-week BM of recipients. **(H)** Donor-derived nucleated reconstitution in PB after the secondary transplantation. Data were shown from one representative experiment, n = 5 in Y-HSC1 and A-EMPP6, n =6 in Y-EMPP6 and A-HSC1. **(I)** Donor-derived myeloid, B and T cell reconstitution in PB after the secondary transplantation. **(J)** Reconstitution of donor-derived B-cell subsets in 16-week BM (left) and SP (right) of recipients after the secondary transplantation. Data were shown as mean ± SD. \**P* ≤ 0.05, \*\**P* ≤ 0.01, \*\*\**P* ≤ 0.001, and \*\*\*\**P* ≤ 0.0001 (multiple t-test for Fig. 6B; one-way ANOVA for Fig. 6D**-J** with following comparisons performed: Y-HSC1 vs. Y-EMPP6, Y-HSC1 vs. A-HSC1, Y-EMPP6 vs. A-EMPP6, A-HSC1 vs. A-EMPP6). See also Fig. S6.

Collectively, these results suggested that EMPP6 exhibited better resilience to age-related stresses than HSC1 in its relatively maintained lymphoid output during ageing. Moreover, the distinct behavior of aged EMPP6 from aged HSC1 suggested that the EMPP6 may exist independently from HSC1 before ageing.

### EMPP6 initiates an alternative lymphoid differentiation pathway likely independent from HSC1

Given the substantial lymphoid output of EMPP6 into ageing, in stark contrast to the nearly indiscernible lymphoid reconstitution of A-HSC1, we hypothesized that EMPP6 might diverge from HSC1 during young adulthood, which was also hinted by trajectory inferring (**Fig. S5D**). Since transplantation serves as a high-stress setting where HSC1 might give rise to EMPP6 due to its plasticity under pressure, we turned to paired daughter cell (PDC) assay in combination with scRNA-seq to investigate their progeny relationship in a much less stressed condition **(Fig. 7A)**. Division dynamics and single cell colony assay results of HSC1 in two different culture conditions, (SCF+TPO (S-T),^40–42^ versus SCF+IL-12 (S-12), verify the technical feasibility and exclude the potential biases introduced by culture conditions **(Fig. S7A-B)**. A random forest classifier was then trained and tested on an integrated dataset comprising homeostatic HSC1, EMPP6, EMPP5, MPP Ly-I and MPP1-4 to elucidate PDC identities (**Fig. 7B, C**). Query data was composed of PDCs as well as parallelly sorted fresh HSC1 and EMPP6 **(Fig. S7C)**. Supporting our assumption, while most PDCs were classified as MPP1 and EMPP5, barely any EMPP6 were found in HSC1-derived PDCs and vice versa **(Fig. 7D, S7D)**. Notably, a higher proportion of EMPP6-derived PDCs were classified as CD135^+^ lymphoid progenitors, i.e., MPP Ly-I and MPP4 **(Fig. 7D)**, indicating that this method could reveal the intrinsic lineage bias of the starting cells. High-importance features for model classification were acquired, in which acknowledged population markers were correctly identified (**Supplemental Table S4**), verifying that the model was able to capture biological characteristics separating these populations. Effect of culture conditions was also ruled out, as no significant difference was observed in the colony-forming efficiency between EMPP6 and EMPP5 **(Fig. S7E)**. Interestingly, S-12 condition supported more symmetric self-renewal events than S-T (**Fig. S7F**), consistent with our previous finding.^28^

**Fig. 7.**
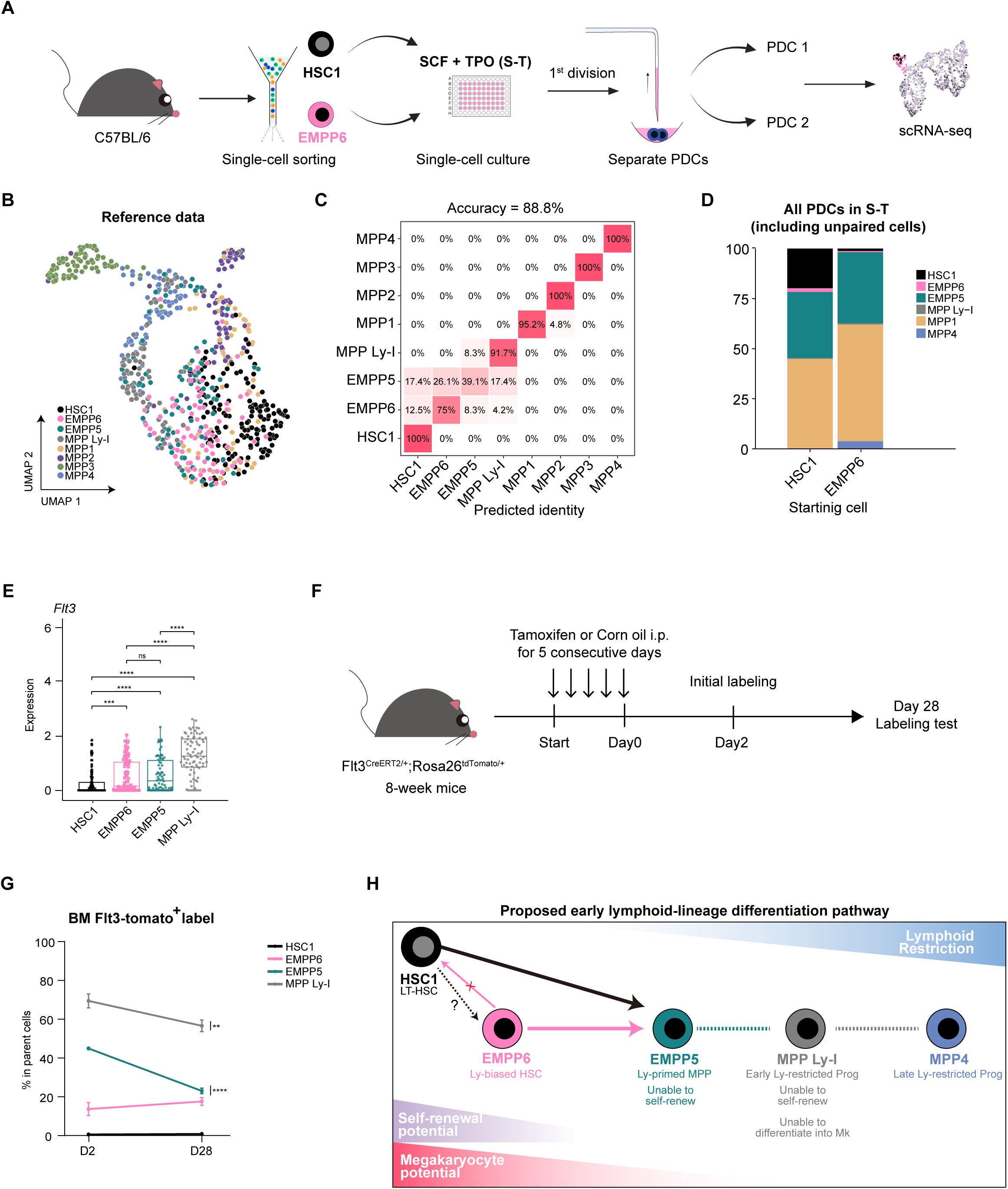
EMPP6 independently initiates a lymphoid differentiation pathway. **(A)** Schematic plot illustrating the experiment design of pair-daughter cell (PDC) assay. See method for details. **(B)** UMAP visualization of integrated data used as reference for random forest classifier. HSC1, 159 cells; EMPP6, 81 cells; EMPP5 78 cells; MPP Ly-I, 80 cells; MPP1, 72 cells; MPP2, 72 cells; MPP3, 76 cells; MPP4, 78 cells. HSC1 cells were from Fig. 2. EMPP6, EMPP5 and MPP Ly-I were from Fig. 5. MPP1-4 were from Dong et al.^57^. **(C)** Confusion matrix calculated by applying the trained classifier to all reference data. **(D)** Stacked bar plot showing classification results for both HSC1-and EMPP6-derived PDCs. All PDCs regardless of whether paired or not were included. **(E)** Expression of *Flt3* in indicated cell populations under homeostasis. **(F)** Workflow for tracing BM HSC1 and EMPP6 under homeostasis using *Flt3*^CreER^/*R26*^tdTomato^ mice. **(G)** Labeling efficiency for each cell population after 4-week tracing in *Flt3*^CreER^/*R26*^tdTomato^ mice. **(H)** Refined model for lymphoid-lineage differentiation pathway within the LSK compartment. Dashed arrows indicate relationships that warrant further functional validation. Data were shown as mean ± SD. \*\**P* ≤ 0.01, \*\*\**P* ≤ 0.001, and \*\*\*\**P* ≤ 0.0001 (wilcoxon test for Fig. 7E; multiple t-test for Fig. 7G) See also Fig. S7 and Supplemental Table S4.

To further illustrate the relationship between HSC1 and EMPP6 in vivo, the *Flt3*^CreER^/*R26*^tdTomato^ mice were used to trace the labeling efficiency of respective cell populations under homeostasis given the differential expression of *Flt3* in EMPP6 than HSC1 (**Fig. 7E**). The initial labeling efficiency at day 2 and after 28-day chasing was compared after 5 consecutive days injection of tamoxifen **(Fig. 7F, S7G-H)**. As shown in **Fig. 7G**, 2-day after tamoxifen induction, there were 13.70%, 44.97%, and 69.43% Tomato^+^ cells within the EMPP6, EMPP5, and MPP Ly-I cells respectively. Whereas there was no Tomato^+^ cells in HSC1 at the initial labeling, which were consistent with the *Flt3* expression patterns among those populations **(Fig. 7E)**. We hypothesized that the labeling efficiency of EMPP6 will be changed after proliferation or differentiation. Four-week later, the Tomato^+^ frequencies of EMPP5 and MPP Ly-I decreased dramatically to 23.07% and 56.63%, respectively, indicating either the supplement from upstream populations (probably from unlabeled EMPP6 or HSC1) or the differentiation towards downstream cells **(Fig. 7G)**. Notably, consistent with the quiescent property of EMPP6 **(Fig. 3B)**, the Tomato^+^ EMPP6 maintained unchanged during 4-week chasing, suggesting that either the low proliferation or rarely differentiation of labelled EMPP6, or both under homeostasis. There was neither Tomato^+^ HSC1 cells after 28 days chasing, suggesting that EMPP6 especially Tomato^+^ EMPP6 rarely generate HSC1 under steady state. Taken together, all those results indicated the independent probability between EMPP6 and HSC1 under both stress and homeostasis conditions.

Stitching all results together, we here propose a refined model for the lymphoid differentiation pathway within the LSK compartment (**Fig. 7H**): Separated from HSC1 at least in the young adulthood, EMPP6, the Ly-biased HSCs initiates an alternative route which converges with the one from HSC1 at the lymphoid-primed EMPP5 and finally reaches the lymphoid-restricted progenitors represented by MPP Ly-I and MPP4.

## DISCUSSION

In this study, we first compared EPCR^+^ (HSC2) and EPCR^−^ (DN) cells in the CD150^−^ fraction. Although approximately 75% HSC2 cells were overlapped with CD45^+^EPCR^+^CD48^−^CD150^−^ cells,^23^ the functional and molecular features of which have been reported, the heterogenous composition of HSC2 and their EPCR^−^ counterparts have not been investigated yet. By assessing the reconstituting ability of DN, we ensured the role of EPCR to purify more primitive HSPCs for both the CD150^+^ or CD150^−^ fraction. By examining the heterogeneity within HSC2, three subsets (EMPP6, EMPP5, and MPP Ly-I) were identified at both functional and molecular levels. Despite previously detailly reported MPP5^36^ and MPP Ly-I^5^, they have never been compared together. By transplantation and scRNA-seq, we uncovered the functional and molecular rankings among the three subsets, and proposed a refined hematopoiesis model where EMPP5 and MPP Ly-I were downstream of EMPP6 in the hierarchy. Although the hierarchy marked by the expression of CD34 and CD135 in LSK cells has previously been established by Yang et al,^43^ EMPP6 was a much purified population based on additional combination of CD150^−^, CD48^−^ and EPCR^+^ and was less overlapped with CD34^−^FLT3^−^LSK (about 11%, **Figure S8A**). Similarly, EMPP6 was also less overlapped with CD150^−^CD34^−^LSK^44^ and CD45^+^EPCR^+^CD48^−^CD150^−^ (**Figure S8B, C**).^23^ Recently, Somuncular et al. reported CD49b^+^ subset in CD150^+^CD48^−^CD34^−^ LSK with a lymphoid bias,^45^ and nearly 75% of EMPP6 were found to be CD49b^+^ (**data not shown**). CD49b may thus be a potential marker for further purification of EMPP6.

EMPP6, the Ly-biased HSCs newly identified in this study, differs from previously reported lymphoid progenitors by its self-renewal and multi-lineage potential. A recent study also described a phenotypically distinct EPCR^+^ MPP6 population (EPCR^+^CD150^−^CD48^−^CD34^−^ FLT3^−^LSK, termed LT-HSC2) with exceptional quiescence and the earliest lymphoid priming.^46^ EMPP6 in our study is phenotypically identical to LT-HSC2, and our data support a similar quiescent state for EMPP6 and HSC1. Functionally, we defined this population as Ly-biased HSCs rather than LT-HSC2 because: (i) despite overall comparable chimerism to HSC1 in bulk serial transplantation (in both studies), its multilineage differentiation ability skewed to lymphoid lineages, as mirrored by the gradual decrease of myeloid and platelet output in our experiments; and (ii) clonal-level serial transplantation revealed inferior self-renewal of EMPP6, with declined lymphoid-restricted output and exhausted myeloid/platelet contribution after serial transplant— consistent with the previously reported functional ST-HSC.^42^ We propose that the truly functional multipotent HSC clones within EMPP6 may be too rare to detect, or that the functional HSC state is too transient at this stage, given the continuum from HSCs to MPPs. We therefore prefer the term Ly-biased HSCs and regard it as the earliest lymphoid priming event in the hematopoietic hierarchy. (**Fig. 7H**).

Notably, in contrast to HSC1, EMPP6 displayed aging-resilient property by maintaining continuous lymphoid-biased reconstitution and supporting ongoing lymphopoiesis (reflected by generation of pro B cells at 16 weeks after transplantation). Similar to our founding, Wang et al. found that CD150^low^ HSC in aged mice exhibited younger features and could alleviate ageing phenotypes,^7^ which was also insinuated by a recent study proposing that the altered ratio of LT-and ST-HSC resulted in myeloid-biased hematopoiesis in ageing.^47^ To combat aging, Non-expanded HSC clones with repopulating capacity was reported during ageing,^6^ and Flk2^+^ MPPs (MPP^F^) was also revealed to protect the downstream progenitor cells from inheriting age-related properties from HSCs.^48^ Given the similarities of EMPP6 shared with these populations, we proposed EMPP6 as also an ageing-resilient population positioned far upstream of MPP^F^, strengthening the notion that aging of HSCs is inconsequential to progenitor cell function. We suppose that there exist at least two possible mechanisms as to why EMPP6 maintains the lymphoid potential during aging. Firstly, in contrast to abnormally expanded HSC1, EMPP6 only mildly expands. Considering its highly quiescent feature suggested by cell cycle analysis, we assume that EMPP6 maintains this highly quiescent state for the lifetime which protects it from proliferation-induced damages. Secondly, EMPP6 exhibited higher score of “q4” gene set^7^. This gene set is upregulated in young CD150^+^HSC compared to its aged counterparts, and enriched for “Regulation of leukocyte cell-cell adhesion”. This is consistent with EMPP6 showing prominent cytoskeleton-related terms. These cytoskeleton-related functions might be, for some reason, well maintained in EMPP6 during ageing and therefore support its targeting towards specific niches, its mobility across BM and different lymphoid organs or its reception of the lymphoid differentiation signals.

Progeny relationship between the CD150^+^ and CD150^−^ fraction has long been a point of controversy. While some classic studies suggested that the CD150^+^ fraction gave rise to the CD150^−^ fraction, recently many studies have suggested independent embryonic origins of these two populations.^49–51^ One of the confounding factors is the purity of CD150^−^ fraction. After highly purifying the CD150^−^ fraction, we were able to exclude interference of EMPP5 and MPP Ly-I which dominate in the CD150^−^ fraction. Through trajectory analysis, PDC assay, and in vivo short-term lineaging tracing after labeling EMPP6, we offered new evidence supporting the strong independent probability of EMPP6 from HSC1 at least in the young adulthood. These pieces of information lead us to a somewhat bold hypothesis: EMPP6, or at least some clones within EMPP6, might be embryonic-originated, persisting into adulthood and contributing to lymphopoiesis, similar to previously reported eMPP.^52,53^ As HSC1 cells could not be labelled and traced in our current model, i.e. the HSC1-derived EMPP6 could not be assessed, HSC1-specific and -efficient labeling model from the embryonic stage would be an appropriate approach to further investigate this.

Collectively, we identified a self-renewable and aging-resilient Ly-biased HSCs, from which, we assumed, an alternatively HSC-independent early-stage lymphoid differentiation pathway was initiated, providing both conceptual and translational insights into hematopoiesis and immunology.

### Limitations of the Study

Our study mainly relied on transplantation settings in which Ly-biased HSCs (EMPP6) and Ly-primed MPPs (EMPP5) were defined. The role of these cells in physiological settings remains to be elucidated utilizing more specific tracing models with durable chasing until aging stage. We assumed the relationship between Ly-biased HSCs (EMPP6) and LT-HSC based on scRNA-seq of PDCs and short-term lineage tracing during young stage. The origin of EMPP6 during embryonic stage and the exact relationship with LT-HSC warrant further investigation using more specific lineage tracing model. In addition, the number of differentially expressed genes between Ly-biased HSCs and LT-HSC was limited. Previous studies have highlighted importance of epigenetic regulation in HSC fate decision.^54,55^ Therefore, epigenetic studies are required to further understand potential mechanisms underlying their functional divergences. Specific relationship between EMPP6 and EMPP5 and MPP1 should be clarified to have a more comprehensive view of myeloid and lymphoid lineage commitments.

## Supporting information

All supplemental figures

## RESOURCE AVAILABILITY

### Materials availability

There are no restrictions to the availability of all materials mentioned in the manuscript.

### Data and code availability

Single-cell RNA-seq data have been deposited at GSE301843 (HSC1 and HSC2 cells), GSE301841 (EMPP6, EMPP5 and MPP Ly-I cells), GSE302376 (HSC1-derived PDCs), and GSE319040 (EMPP6-derived PDCs), and are publicly available as of the date of publication. Mass spectrometry data was supplemented in **Supplemental Table S6**. All original code has been deposited at https://github.com/Mayao000/Ly-HSCs and is publicly available as of the date of publication. For any additional information, please contact.

## ACKNOWLEDGMENTS

We thank Toshio Suda for his critical reading of our paper, Ping Zhu for his help in bioinformatics training, Jianlong Sun for his advice on lineage tracing experiment, Bingqian Chu, Mingming Niu, and Hong Wang for their generous help in performing mass spectrometry and support for related bioinformatic analyses, and our laboratory members for their constructive suggestions.

## AUTHOR CONTRIBUTIONS

YM designed and performed experiments, analyzed data, performed bioinformatic analysis, and wrote the manuscript. XL performed experiments and wrote the manuscript. XL, LW, YX and JZ assisted in the experiments. JW assisted in the experiments and funding application. B Z, and Y L assisted in the project design and lineage tracing experiment. TC, HE and FD conceived and supervised the project, acquired funding, and wrote the manuscript.

## Supplemental Figure Legends

**Fig. S1. Bone marrow and spleen analyses 16 weeks after transplantation, Related to** Fig. 1**. (A)** Representative flow cytometry showing simultaneous GFP and CD41 expression in donor-derived cells.

**(B)** Representative flow cytometry gating of donor-derived myeloid, B and T cells.

**(C)** Representative flow cytometry gating of donor proportion in PB platelets.

**(D)** Donor-derived reconstitution in the 16-week bone marrow (BM). Data were shown from one representative experiment; HSC1, n = 7; HSC2, n = 3; DN, n = 7.

**(E)** Donor-derived reconstitution in the 16-week spleen (SP).

**(F)** Representative High Content Analysis photos (5x) of colonies formed by single HSC1 cells (left) and HSC2 cells (right).

**(G)** Colony-forming efficiency of HSC1 and HSC2 cells. Data were collected from four independent experiments.

**(H)** Proportion of colony types formed by HSC1 and HSC2. nm, neutrophils and/or macrophages; nmE/Mk, nm with erythroblasts and/or megakaryocytes.

**(I)** Donor-derived nucleated and mature-lineage reconstitution at 4, 8, 12, and 16 weeks after transplantation with single HSC1 (top) and HSC2 (bottom) cells. HSC1, n = 25; HSC2, n = 19.

**(J)** Representative flow cytometry profile illustrating the calculation of L/M ratio. Data were shown as mean ± SD. \**P* ≤ 0.05, \*\**P* ≤ 0.01, \*\*\**P* ≤ 0.001, and \*\*\*\**P* ≤ 0.0001 (one-way ANOVA for **Fig. S1D, E**; unpaired Student’s t-test for **Fig. S1 G, H**).

**Fig. S2. HSC2 cells show varying levels of CD34 and CD135 expression. Related to** Fig. 2**. (A)** UMAP visualization (left) of 6 clusters identified in HSC1 and HSC2 data together with data of Sommerkamp et al. and dot plot (right) showing immunophenotypic HSCs and MPPs scoring for each cluster.

**(B)** Markers of each cluster in **Fig. S2A**.

**(C)** Representative flow cytometry profiles (left) and quantitative analysis (right) of EPCR expression in traditional HSC (tradHSC) and MPP1. Colors in the stacked bar plot represented overlapped populations.

**(D)** Representative flow cytometry profiles (left) and quantitative analysis (right) of EPCR expression in MPP5-7. Colors in the stacked bar plot represented overlapped populations.

**(E)** MFI of EPCR in HSC1, HSC2 and MPP1-7 was shown. Data were shown as mean ± SD. \**P* ≤ 0.05, \*\**P* ≤ 0.01, \*\*\**P* ≤ 0.001, and \*\*\*\**P* ≤ 0.0001 (one-way ANOVA for **Fig. S2E**).

**Fig. S3. Functional assessment of HSC2 subsets. Related to** Fig. 3**. (A)** Statistical analyses of donor-derived lineage reconstitution in PB. Data were collected from three independent experiments, HSC1, n = 16; EMPP6, n = 16; EMPP5, n = 16; MPP Ly-I, n = 15.

**(B)** Donor-derived reconstitution in the 16-week BM of primary recipients. Data were collected from 3 independent experiments, HSC1, n = 11; EMPP6, n = 11; EMPP5, n = 11; MPP Ly-I, n = 8.

**(C)** Donor-derived reconstitution in the 16-week SP of primary recipients.

**(D)** Representative flow cytometry gating of D-tradHSC, D-EMPP6, D-EMPP5, D-MPP Ly-I, D-MPP1, D-MPP2, D-MPP3 and D-MPP4.

**(E)** Representative flow cytometry gating of D-CLP.

**(F)** Representative flow cytometry gating of D-CMP, D-GMP, D-MEP and D-MkP.

**(G)** Donor-derived hematopoietic stem and progenitor cells (HSPC) in the 16-week BM of primary recipients. Data were collected from two independent experiments, n = 6 for each group.

**(H)** Donor-derived HSPCs in the 16-week BM of secondary recipients. Data were collected from three independent experiments, n = 13 for each group. Data were shown as mean ± SD. \**P* ≤ 0.05, \*\**P* ≤ 0.01, \*\*\**P* ≤ 0.001, and \*\*\*\**P* ≤ 0.0001 (one-way ANOVA for **Fig. S3A-C**; only HSC1 and EMPP6 were compared using unpaired Student’s t-test in **Fig. S3G**; unpaired Student’s t-test for **Fig. S3H**).

**Fig. S4. Clonal types of HSC1 and EMPP6 cells. Related to** Fig. 4**. (A)** Dot plot showing lineage reconstitution levels in each reconstituted mouse by single HSC1 and EMPP6 cells 4, 8, 12, and 16 weeks after transplantation. Colors filling dots (blue) represent reconstitution level. Numbers in dots indicate specific reconstitution level. Colors lining dots represent clonal types. HSC1, n = 24; EMPP6, n = 28.

**(B)** Proportion of all reconstituted mice displayed in **Fig. S4B** (outer) and clone types (inner) at 4, 8, 12 and 16 weeks.

**Fig. S5 Distinct transcriptomic profiles of EMPP6, EMPP5 and MPP Ly-1. Related to** Fig. 5**. (A)** UMAP visualization of 81 EMPP6, 78 EMPP5, and 80 MPP Ly-I cells, collected from 6 mice.

**(B)** Results of GESA using the “hallmark gene sets” from MsigDB between indicated subpopulations.

**(C)** Radar plot showing enrichment scores of EMPP6, EMPP5 and MPP Ly-1 based on cell-type gene sets.^5^

**(D)** 3D visualization of inferred trajectory by diffusion map of HSC1, EMPP6, EMPP5, MPP Ly-I, MPP1, MPP2, MPP3, MPP4, CMP, CLP, GMP and MEP cells.

**(E-F)** Enrichment scoring using HSC-related^33–35^ gene sets. \**P* ≤ 0.05, \*\**P* ≤ 0.01, \*\*\**P* ≤ 0.001, and \*\*\*\**P* ≤ 0.0001 (wilcoxon test).

**Fig. S6. EMPP6 was of higher functional fitness than HSC1 in dealing with age-related stress. Related to** Fig. 6**. (A-B)** Donor-derived nucleated and myeloid cell reconstitution in the 16-week BM (**A**) and 16-week SP (**B**) of primary recipients. Data were shown from one representative experiment, n = 6 in both groups.

**(C)** Representative gating of BM and SP B-cell subsets.

**(D)** Representative gating of BM and SP CD4^+^T-cell subsets.

**(E)** Representative gating of BM and SP CD8^+^T-cell subsets.

**(F)** Donor-derived CD4^+^T-cell (left) and CD8^+^T-cell (right) subsets in the 16-week SP of primary recipients.

**(G)** Donor-derived HSPC reconstitution in the 16-week BM of primary recipients. Data were shown as mean ± SD. \**P* ≤ 0.05, \*\**P* ≤ 0.01, \*\*\**P* ≤ 0.001, and \*\*\*\**P* ≤ 0.0001 (one-way ANOVA with following comparisons performed: Y-HSC1 vs. Y-EMPP6, Y-HSC1 vs. A-HSC1, Y-EMPP6 vs. A-EMPP6, A-HSC1 vs. A-EMPP6).

**Fig. S7. Validation of pair-daughter cell assay. Related to** Fig. 7**. (A)** Division kinetics of HSC1 cells cultured with SCF+TPO (S-T) and SCF+IL-12 (S-12) and EMPP6 cells cultured with S-T. When 2 cells/well were seen, single starting cells were considered to have divided once; 3 or 4 cells/well, considered to have divided twice; ≥5 cells/well, considered to have divided three times or more.

**(B)** Proportion of colony types formed by HSC1 in S-T and S-12 conditions. nm, neutrophils and/or macrophages; nmE/Mk, nm with erythroblasts and/or megakaryocytes.

**(C)** UMAP visualization of the query data for the classifier. HSC1 Fresh, 42 cells; EMPP6 Fresh, 40 cells; HSC1 S-T PDCs, 267 cells; HSC1 S-12 PDCs, 331 cells; EMPP6 S-T PDCs, 319 cells.

**(D)** Stacked bar plot showing PDCs classification results in S-T and S-12. All PDCs regardless of whether paired or not were included.

**(E)** Single-cell colony assay of HSC1, EMPP6, EMPP5, MPP Ly-I, MPP3, and MPP4. The percentage of colony-forming efficiency is shown.

**(F)** Pie chart showing HSC1 division types in S-T condition. Only paired data were included. SS, symmetric self-renewal; AS, asymmetric self-renewal; SD, symmetric differentiation.

**(G-H)** Representative flow cytometry gating of each cell population in *Flt3*^CreER^/*R26*^tdTomato^ mice at Day 2 (**G**) and Day 28 (**H**) after corn oil or tamoxifen injection. Data were shown as mean ± SD. \**P* ≤ 0.05, \*\**P* ≤ 0.01, \*\*\**P* ≤ 0.001, and \*\*\*\**P* ≤ 0.0001 (one way ANOVA for **Fig. S7E**).

## STAR Methods

### KEY RESOURCES TABLE

| REAGENT or RESOURCE | SOURCE | IDENTIFIER |
|---|---|---|
| Antibodies |  |  |
| CD127 PE (clone A7R34) | eBioscience | Cat# 12-1271-81; RRID: AB_465843 |
| CD127 PE-Cy7 (clone A7R34) | BioLegend | Cat# 135014; RRID: AB_1937265 |
| CD135 PE (clone A2F10) | BioLegend | Cat# 135306; RRID: AB_1877217 |
| CD135 APC (clone A2F10) | BioLegend | Cat# 135310; RRID: AB_2107050 |
| CD150 PE (clone TC15-12F12.2) | BioLegend | Cat# 115904; RRID: AB_313683 |
| CD150 APC (clone TC15-12F12.2) | BioLegend | Cat# 115909; RRID: AB_493461 |
| CD150 BV421 (clone TC15-12F12.2) | BioLegend | Cat# 115926; RRID: AB_2562190 |
| CD16/32 PerCP-eFluor710 (clone 93) | eBioscience | Cat# 46-0161-82; RRID: AB_2016701 |
| CD19 PE (clone 1D3/CD19) | BioLegend | Cat# 152408; RRID: AB_2629817 |
| CD21 FITC (clone 7E9) | BioLegend | Cat# 123407; RRID:AB_940403 |
| CD23 PerCP-Cy5.5 (clone B3B4) | Invitrogen | Cat# 46-0232-82; RRID: AB_2573674 |
| CD3e APC-eFluor780 (clone 145-2C11) | eBioscience | Cat# 47-0031-82; RRID: AB_11149861 |
| CD3e PE (clone 145-2C11) | Invitrogen | Cat# 12-0031-82; RRID: AB_465496 |
| CD3e PE-Cy7 (clone 145-2C11) | eBioscience | Cat# 25-0031-81; RRID: AB_469571 |
| CD34 Biotin (clone RAM34) | eBioscience | Cat# 13-0341-82; RRID: AB_466425 |
| CD34 FITC (clone RAM34) | eBioscience | Cat# 11-0341-82; RRID: AB_465021 |
| CD4 APC-Cy7 (clone GK1.5) | BioLegend | Cat# 100414; RRID: AB_312699 |
| CD4 PE-Cy7 (clone GK1.5) | BioLegend | Cat# 100422; RRID: AB_312707 |
| CD41a APC (clone eBioMWReg30 (MWReg30)) | eBioscience | Cat# 17-0411-82; RRID: AB_1603237 |
| CD43 APC (clone S7) | BD Biosciences | Cat# 560663; RRID: AB_1727479 |
| CD43 PE-Cy7 (clone S11) | BioLegend | Cat# 143209; RRID: AB_2564348 |
| CD44 APC (clone IM7) | Invitrogen | Cat# 17-0441-82; RRID: AB_469390 |
| CD45.2 FITC (clone 104) | BioLegend | Cat# 109806; RRID:AB_313443 |
| CD45.2 PE (clone 104) | BioLegend | Cat# 109808; RRID:AB_313445 |
| CD45.2 PE-Cy7 (clone 104) | BioLegend | Cat# 109830; RRID:AB_1186098 |
| CD48 AF700 (clone HM48-1) | BioLegend | Cat# 103426; RRID: AB_10612755 |
| CD48 PE-Cy7 (clone HM48-1) | BioLegend | Cat# 103424; RRID: AB_2075049 |
| CD5 PE (clone 53-7.3) | Invitrogen | Cat# 12-0051-82; RRID: AB_465523 |
| CD62L FITC (clone MEL-14) | BioLegend | Cat# 104406; RRID:AB_313093 |
| CD8a APC (clone 53-6.7) | BioLegend | Cat# 100712; RRID: AB_312751 |
| CD8a APC-eFluor780 (clone 53-6.7) | eBioscience | Cat# 47-0081-82; RRID: AB_1272185 |
| CD8a PE-Cy7 (clone 53-6.7) | BioLegend | Cat# 100722; RRID: AB_312761 |
| CD8a PerCP-Cy5.5 (clone 53-6.7) | BioLegend | Cat# 100734; RRID:AB_2075238 |
| B220 APC-Cy7 (clone RA3-6B2) | BioLegend | Cat# 103224; RRID: AB_313007 |
| B220 PE-Cy7 (clone RA3-6B2) | eBioscience | Cat# 25-0452-82; RRID: AB_469627 |
| B220 PerCP-Cy5.5 (clone RA3-6B2) | BioLegend | Cat# 103236; RRID: AB_893354 |
| c-Kit BV605 (clone 2B8) | BioLegend | Cat# 105847; RRID: AB_2783047 |
| EPCR PerCP-Cy5.5 (clone eBio1560 (1560)) | eBioscience | Cat# 46-2012-80; RRID: AB_10718383 |
| IgM APC (clone RMM-1) | BioLegend | Cat# 406509; RRID:AB_315059 |
| Mac-1 APC-Cy7 (clone M1/70) | BioLegend | Cat# 101226; RRID: AB_830642 |
| Mac-1 PE-Cy7 (clone M1/70) | BioLegend | Cat# 101216; RRID: AB_312799 |
| NK1.1 PE-Cy7 (clone PK136) | Invitrogen | Cat# 25-5941-82; RRID: AB_469665 |
| Sca-1 AF700 (clone D7) | eBioscience | Cat# 56-5981-82; RRID: AB_657836 |
| Sca-1 BV785 (clone D7) | BioLegend | Cat# 108139; RRID: AB_2565957 |
| Gr-1 APC-Cy7 (clone RB6-8C5) | BioLegend | Cat# 108424; RRID: AB_2137485 |
| Gr-1 PE-Cy7 (clone RB6-8C5) | eBioscience | Cat# 25-5931-82; RRID: AB_469663 |
| Streptavidin APC-Cy7 | BioLegend | Cat# 405208; RRID: AB_N/A |
| Ter-119 APC-Cy7 (clone TER-119) | BioLegend | Cat# 116223; RRID: AB_2137788 |
| Ter-119 PE-Cy7 (clone TER-119) | BioLegend | Cat# 116222; RRID: AB_2281408 |
| Chemicals, peptides, and recombinant proteins |  |  |
| Recombinant Murine SCF | Peprotech | Cat# AF-250-03 |
| Recombinant Murine TPO | Peprotech | Cat# AF-315-14 |
| Recombinant Murine IL-3 | Peprotech | Cat# AF-213-13 |
| Recombinant Murine IL-12 p70 | Peprotech | Cat# 210-12 |
| Polyvinyl alcohol (PVA) | Sigma-Aldrich | Cat# 363081 |
| Critical commercial assays |  |  |
| CD117 MicroBeads, mouse | Miltenyi Biotec | Cat# 130-091-224 |
| LS Columns | Miltenyi Biotec | Cat# 130-042-401 |
| MS Columns | Miltenyi Biotec | Cat# 130-042-201 |
| AMPure XP beads | Beckman | Cat# A63882 |
| Dynabeads Myone streptavidin C1 beads | Invitrogen | Cat# 65001 |
| Kapa Hyper Prep Kit | KAPA | Cat# KK8504 |
| Bioanalyzer high sensitivity DNA chip | Agilent | Cat# 5067-4626 |
| Qubit dsDNA assay kit | Thermo Fisher | Cat# Q32854 |
| Deposited data |  |  |
| Raw scRNA-seq data | This paper | GSE301843 (HSC1 and HSC2 cells), GSE301841 (EMPP6, EMPP5 and MPP Ly-I cells), GSE302376 (HSC1-derived PDCs), and GSE319040 (EMPP6-derived PDCs) |
| Experimental models: Organisms/strains |  |  |
| Mouse: C57BL/6 (CD45.2) | Beijing Vital River Laboratory Animal Technology | N/A |
| Mouse: B6.SJL (CD45.1) | Beijing Vital River Laboratory Animal Technology | N/A |
| Mouse: β-actin-GFP transgenic reporter mice (C57BL/6-CD45.2 background) | In house breeding | N/A |
| Mouse: B6.Cg-Tg(FucciG1)#610Bsi (mKO2-hCdt1(30/120)) | RIKEN BRC, Japan | N/A |
| Mouse: B6.Cg-Tg(FucciS/G2/M)#474Bsi (mAG-hGeminin(1/110)) | RIKEN BRC, Japan | N/A |
| Software and algorithms |  |  |
| FlowJo v.10.0.7 | BD Bioscience | https://www.flowjo.com/ |
| Prism v.8.0.2 | GraphPad | https://www.graphpad.com/scientificsoftware/prism/ |
| Microsoft Excel | Microsoft | https://www.microsoft.com/en-gb/ |
| Python v.3.6.15 | Anaconda, Inc | https://docs.conda.io/en/latest/miniconda.html |
| R v.4.2.2 | R core team | https://www.r-project.org/ |
| Trimmomatic v.0.39 | Bolger et al.<sup>58</sup> | http://www.usadellab.org/cms/?page=trimmomatic |
| HISAT2 v.2.2.1 | Kim et al.<sup>59</sup> | http://daehwankimlab.github.io/hisat2/ |
| HTSeq v.0.11.3 | Putri et al.<sup>60</sup> | https://htseq.readthedocs.io/en/latest/ |
| Cell Ranger v.3.0 | 10x Genomics | https://support.10xgenomics.com/single-cellgene-expression/software/downloads/latest |
| R Seurat v.5.0.0 | Stuart et al.<sup>61</sup> | https://satijalab.org/seurat/ |
| R clusterProfiler v.4.6.2 | Wu et al.<sup>62</sup> | https://bioconductor.org/packages/release/bioc/html/clusterProfiler.html |
| R org.Mm.eg.db v.3.16.0 | Bioconductor core team | https://bioconductor.org/packages/release/data/annotation/html/org.Mm.eg.db.html |
| R monocle3 v.1.3.4 | Cao et al.<sup>63</sup> | https://cole-trapnell-lab.github.io/monocle3/ |
| Python Scanpy v.1.7.2 | Wolf et al.<sup>64</sup> | https://scanpy.readthedocs.io/en/stable/ |
| R reticulate v.1.34.0 | Ushey et al.<sup>65</sup> | https://rstudio.github.io/reticulate/ |
| R plotly v.4.10.3 | Sievert<sup>66</sup> | https://plotly.com/r/ |
| R caret v.6.0.94 | Kuhn<sup>67</sup> | https://github.com/topepo/caret |
| R randomForest v.4.7.1.2 | Liaw et al.<sup>68</sup> | https://cran.r-project.org/web/packages/randomForest/index.html |

### EXPERIMENTAL MODEL AND STUDY PARTICIPANT DETAILS

#### Animals

For in vitro culture and single-cell RNA sequencing (scRNA-seq) experiments, 8 ∼ 10-week-old female C57BL/6 (CD45.2) mice were used. The mice were purchased from Beijing Vital River Laboratory Animal Technology. For transplantation experiments, 8 ∼ 10-week-old female β-actin–GFP transgenic mice (CD45.2) bred in the animal facility of the Institute of Hematology were used as donors, and 8 ∼ 10-week-old female C57BL/6 mice were used as recipients. For cell cycle analyses, B6.Cg-Tg(FucciG1) #610Bsi (mKO2-hCdt1(30/120)) and B6.Cg-Tg(FucciS/G2/M) #474Bsi (Fucci-mAG, CD45.2)^38^ were first obtained from RIKEN BRC, Japan, and were then intercrossed with B6.SJL mice to acquire Fucci mice with a CD45.1 background.^26^ Fucci dual reporter mice (Fucci) were generated through mating male Fucci-mKO2-mAG (CD45.1) with female Fucci-mAG (CD45.1) mice, and 8 ∼ 10-week-old female Fucci littermates were used for experiments. For ageing-related functional assay, 91-week old female C57BL/6 (CD45.2) mice were used as donors, and 8 ∼ 10-week-old female B6.SJL (CD45.1) mice were used as recipients. The abovementioned mice were maintained in the animal facility of the Institute of Hematology with strict 12-h light-dark cycles to ensure regular circadian rhythm, and sterilized water and food were continuously provided. All animal experiments were performed in compliance with the animal care guidelines approved by the Institutional Animal Care and Use Committees of the State Key Laboratory of Experimental Hematology. Efforts were made to reduce animal number and suffering.

### METHOD DETAILS

#### Flow cytometric analysis and sorting

Antibodies used in this study were listed in the **Key Resources Table.**

Peripheral blood (PB), bone marrow (BM) and spleen (SP) samples for analysis of the myeloid/lymphoid lineages were depleted with erythrocytes, stained with corresponding antibodies (see **Supplemental Table S1**) for 30 minutes at 4 in the dark, washed with EDTA-containing PBS (PBE), and resuspended in DAPI-containing PBE. PB samples for analysis of platelets were directly stained with antibodies for 30 minutes at the room temperature in the dark, washed with PBE, and resuspended in DAPI-free PBE. For BM analyses of hematopoietic stem and progenitor cells (HSPCs), whole BM cells (wBMCs) were depleted with erythrocytes and stained with antibodies for 30 (without CD34 staining) or 60 (direct or biotinylated CD34 staining) minutes at 4 in the dark. Lineage markers contained Ter-119, B220 and Gr-1 for the hematopoietic stem cells (HSCs) and multipotent progenitors (MPPs) panel, and contained Ter-119, B220, Gr-1, Mac-1, CD3, CD4 and CD8 for other panels. The analysis was performed on a BD LSRFortessa or FACSAria III.

For flow cytometric sorting of indicated cell populations, wBMCs were enriched using mouse CD117 (c-Kit) MicroBeads purchased from Miltenyi Biotec, and stained with corresponding antibodies. For single-cell experiments, cells were sorted using “single-cell” mode. Otherwise “4-way purity” mode was applied. The sorting was performed on a BD FACSAria III.

#### Transplantation assays

For population transplantation in **Fig. 1B**, **1C** and **Fig. 3**, indicated populations (200 cells each) isolated from 8 ∼ 10-week-old female β-actin–GFP transgenic reporter mice were intravenously injected into lethally irradiated (4.50 Gy twice, at least 4 hrs apart) 8 ∼ 10-week-old female C57BL/6 mice, together with 5 × 10^5^ wBMCs as competitor cells from C57BL/6 mice. PB was collected from the tail veins at indicated weeks after transplantation, and separated into two tubes for analyses of the donor-derived (GFP^+^) myeloid/lymphoid lineages and platelets respectively. For serial transplantation experiments, wBMCs were obtained from primary recipients, mixed and counted within the same group. 1 × 10^7^ cells (per recipient) were then intravenously injected into lethally irradiated 8 ∼ 10-week-old female C57BL/6 mice (secondary recipients). PB was collected at 4, 8, 12 and 16 weeks after transplantation. Same procedures were repeated in the tertiary transplantation.

For ageing-related transplantation in **Fig. 6**, 100 HSC1 and EMPP6 cells were either isolated from 8 ∼ 10-week-old or 91-week old female C57BL/6 (CD45.2) mice and intravenously injected into lethally irradiated 8 ∼ 10-week-old female B6.SJL (CD45.1) mice, together with 5 × 10^5^ wBMCs as competitor cells from B6.SJL mice. PB was collected from the tail veins at 4, 8, 12 and 16 weeks after transplantation. Serial transplantation experiments were performed similarly to the abovementioned.

For single-cell transplantation, single cells were sorted from the indicated populations from 8 ∼ 10-week-old female β-actin–GFP transgenic reporter mice into 96-well plates, mixed with 5 × 10^5^ wBMCs (obtained from C57BL/6 mice) per well, and intravenously injected into lethally irradiated 8 ∼ 10-week-old female C57BL/6 mice. PB was collected at 4, 8, 12 and 16 weeks after transplantation, and separated into two tubes for analyses of the GFP^+^ myeloid/lymphoid lineages and platelets respectively. Cutoff for reconstituted recipients was % nucleated for **Fig. 1** or % (nucleated + platelet) for **Fig. 4** ≥0.1% at any time point. Cutoff for long-term (LT) reconstituted recipients was % nucleated for **Fig. 1** or % (nucleated + platelet) for **Fig. 4** ≥ 0.1% at 16 weeks. For serial transplantation experiments in **Fig. 4**, 2 × 10^7^ wBMCs were obtained from each primary recipients showing ≥1% (nucleated + platelet) reconstitution and one-to-one transplantation was then performed into lethally irradiated 8 ∼ 10-week-old female C57BL/6 mice (secondary recipients). PB was collected at 4, 8, 12 and 16 weeks after transplantation.

#### Single-cell colony assay

Single HSC1 and HSC2 cells were sorted from 8 ∼ 10-week-old female C57BL/6 mice by FACSAria III into U-bottom 96-well plates containing 0.2 ml α-MEM (Gibco) supplemented with 10% FBS (Gibco), 1% penicillin/streptomycin/glutamine (Gibco), 2 mM L-Glutamine (Gibco), 10 ng/ml recombinant murine SCF (Peprotech), 10 ng/ml recombinant murine mTPO (Peprotech), 10 ng/ml recombinant murine IL-3 (Peprotech), and 1 IU/ml human EPO (local hospital) as previously described.^69^ Single HSC1, EMPP6, EMPP5, MPP Ly-I, MPP3 and MPP4 cells were sorted into serum-free F12 medium (Gibco) supplemented with 1% ITS-X (Gibco), 10 mM HEPES (Gibco), 10 μM MEM nonessential amino acids (Gibco), 1% penicillin/streptomycin/glutamine (Gibco), 2mM L-glutamine, 0.055mM 2-mercaptoethanol (Gibco), 1 mg/ml PVA (Sigma-Aldrich), 50 ng/ml mSCF (Peprotech) and 50 ng/ml mTPO (Peprotech) as previously described.^26,28,70^ Cells were cultured for 14 days at 37°C with 5% CO2. For determining colony types, colonies were processed with cytospin and Giemsa staining, and were next morphologically identified and quantitively summarized.

#### Paired daughter cell assay

Single HSC1 and EMPP6 cells were sorted from 8 ∼ 10-week-old female C57BL/6 mice by FACSAria III into U-bottom 96-well plates containing 0.2 ml serum-free F12 medium (Gibco) supplemented with 1% ITS-X (Gibco), 10 mM HEPES (Gibco), 10 μM MEM nonessential amino acids (Gibco), 1% penicillin/streptomycin/glutamine (Gibco), 2mM L-glutamine, 0.055mM 2-mercaptoethanol (Gibco), 1 mg/ml PVA (Sigma-Aldrich), 50 ng/ml mSCF (Peprotech) and 50 ng/ml mTPO (Peprotech) (SCF+TPO, ST) or 50 ng/ml recombinant murine IL-12 (Peprotech) (SCF+IL-12, S12). HSC1 cells were cultured for 48 hours under ST condition and 72 hours under S12 condition, while EMPP6 cells were cultured for 48 hours under ST, when they endured the first division. Individual pair-daughter cells (PDCs) were then picked and separated by a micromanipulator (Narishige, Japan), and were transferred either into aforementioned α-MEM culture medium for single-cell colony assays, or into single-cell lysis buffer for single-cell transcriptome library preparation and sequencing.

#### Preparation of scRNA-seq and mass spectrometry data

Accession to all sequencing data was listed in the **Key Resources Table**.

##### Preparation of scRNA-seq library

Single cells were either sorted by flow cytometry or manually transferred by micromanipulator in the case of PDCs into individual 0.2 ml 8-strip PCR tubes containing 2.25 µL lysis buffer. First-strand cDNAs were synthesized using 25-nucleotide oligo-dT barcoding primers and template switch oligo primers. Following reverse transcription, the barcoded cDNAs were amplified via 20 cycles of PCR. Next, the cDNAs of each cell were pooled together and purified twice using 0.8 × Agencount AMPure XP beads. Biotin tags and indexing sequences were introduced by an additional 4 cycles of PCR amplification using biotinylated primers. PCR products were purified and sonicated (∼ 300 bp by Covaris M220 system). The 3’ ends of the fragmented cDNA fragments were enriched with Dynabeads Myone streptavidin C1 beads (Invitrogen). Finally, the cDNA libraries were generated using a Kapa Hyper Prep Kit (Kapa Biosystems). Quality of libraries were checked using Qubit dsDNA assay kit (Thermo Fisher) and Bioanalyzer high sensitivity DNA chip (Agilent) before sequencing on an Illumina HiSeq platform in paired-end mode (2 × 150 bp) at Novogene (Tianjin, China).

##### Preparation of mass spectrometry data

Indicated populations were sorted and directly deposited into 10 μL of urea-based lysis buffer consisting of 2 M urea, 10% sodium deoxycholate (NaDOC), and 1 M HEPES. Cell suspensions were subjected to repeated cycles of vigorous vortexing for 1 minute followed by 1 minute incubation on ice. Proteolytic digestion was carried out using a combined enzyme system composed of Lys-C (Lysyl Endopeptidase, MS Grade, FUJIFILM Wako, Cat# 125-05061) and sequencing-grade modified trypsin (Promega, Cat# V5113) mixed at a 1:2 ratio. Enzymes were added at a 1:1 enzyme-to-protein ratio, and digestion was performed at 37 °C for 12 hours under continuous agitation at 1000 rpm. Peptide mixtures were then acidified with trifluoroacetic acid (TFA) and clarified by centrifugation at 17,000 × g for 5 minutes. Peptide desalting was performed using Pierce™ C18 Spin Tips (Thermo Scientific, Cat# 84850), which were conditioned with 30 μL of 100% methanol, equilibrated with 30 μL of 0.1% TFA, and peptides were eluted twice using 10 μL of 60% acetonitrile containing 0.1% TFA. Eluted peptides were vacuum-dried and stored at −80 °C. Prior to LC–MS/MS analysis, samples were reconstituted in 0.1% formic acid.

Peptide samples were analyzed on a nanoLC–MS/MS platform composed of a Vanquish Flex UHPLC system (Thermo Fisher Scientific) coupled to an Orbitrap Exploris 480 mass spectrometer (Thermo Fisher Scientific) equipped with a nano-electrospray ionization (NSI) source. Peptide separation was performed on a 15 cm reversed-phase C18 analytical column (ionOpticks; 75 μm inner diameter, 1.7 μm particle size). The mobile phase system consisted of 0.1% formic acid in water (solvent A) and 0.1% formic acid in 80% acetonitrile (solvent B). Ion transfer was achieved through a FAIMS Pro interface operating in standard resolution mode with a compensation voltage of −45 V.

Data acquisition was performed in data-independent acquisition (DIA) mode. MS1 scans were acquired in the Orbitrap analyzer over an m/z range of 350–1250 at a resolution of 120,000, using a maximum injection time of 45 milliseconds and an AGC target equivalent to 3.5 × 10 ions. MS2 spectra were collected at a resolution of 30,000 with a maximum injection time of 54 milliseconds and an AGC target of 1.2 × 10, using higher-energy collisional dissociation (HCD) with a normalized collision energy of 28%. DIA acquisition employed automatically defined isolation windows spanning the precursor mass range of 350–1200 with a 1 m/z overlap, implemented as a scheme of 13 sequential isolation windows.

#### Bioinformatic analysis of transcriptome data

##### Raw data preprocessing

Raw sequencing reads were demultiplexed and assigned to each single cell according to the unique barcode sequences pooled in the sequencing libraries. Poly-A tails and artificial sequences introduced during sequencing library preparation, including template switching oligos (TSOs) and low-quality bases from the 5′ or 3′ ends of each paired reads, were removed by Perl script or Trimmomatic^58^ (v 0.39). Unique molecular identifiers (UMIs) were stored for downstream calculation of gene expression. Sequencing reads passing quality control were aligned to the mouse reference genome (genome version: mm10) using HISAT2^59^ (v 2.2.1) with default settings. The HTSeq package^60^ (v 0.11.3) was used to summarize the uniquely aligned reads for each gene, and duplicated transcripts were removed based on UMIs. The number of transcripts for each gene was calculated as the number of different aligned UMIs. Low-quality cells were removed based on distribution of the number of features, total transcripts detected per cell, and mitochondrial RNA percentage using R package Seurat^61^ (v. 5.0.0). Details were shown in **Supplemental Table S5**.

##### Normalization, dimension reduction and integration

The following procedures were performed with Seurat. Raw counts were normalized using *NormalizeData*. Highly variable features were identified by *FindVariableFeatures* (*nfeature* was determined manually) and scaled by *ScaleData* regressing out cell cycle scores and mitochondrial RNA percentage simultaneously. For dimension reduction and visualization, principal component analysis (PCA) was initially performed using *RunPCA* and then uniform manifold approximation and projection (UMAP) for visualization using *RunUMAP*. For data integration, *IntegrateLayers* was used with the integration method set to CCAIntegration or HarmonyIntegration.

##### Differential expression and enrichment analysis

The following procedures were performed with Seurat. *FindAllMarkers* was used to identify markers for each cluster, whereas *FindMarkers* was used to identify differentially expressed genes (DEGs) between two specified groups. DEGs were those with adjusted *P* value < 0.05. Gene ontology (GO) enrichment analysis was performed by R package clusterProfiler^71^ (v 4.6.2). Terms with adjusted *P* value < 0.05 were considered significant. Gene set enrichment analysis (GSEA) was also performed using clusterProfiler, where input data were genes pre-ranked by log_2_-transformed fold change between two specified groups.

##### Trajectory inferring

R package “monocle3”^63^ (v 1.3.4) was performed on batch-adjusted (by *HarmonyIntegration*) UMAP. Diffusion map analysis was performed using the “Scanpy”^64^ Python package (v 1.7.2) via the “reticulate”^65^ (v 1.34.0) interface in R. Three-dimensional visualization was performed by R package “plotly”^66^ (v 4.10.3).

##### Enrichment scoring

For enrichment scoring in **Fig. S2B** and **S5C**, normalized expression of each feature in a signature gene set was first averaged across all cells within each cell population to obtain an average expression. Then, the average expression values of all features in a signature gene set was summed up and divided by the number of features to generate the final score. For signature scoring in **Fig. 5D and E**, *AddModuleScore* from Seurat was used to calculate score for individual cells.

##### Random forest classification

Raw expression data from published STRT-seq data of MPP1-4 cells^57^ were obtained, processed, and finally integrated with HSC1, EMPP6, EMPP5 and MPP Ly-I to generate the training data for the random forest classifier, which was constructed by R package randomForest (v 4.7.1.2). Briefly, training and test partitions were acquired using *createDataPartition* from R package caret^67^ (v 6.0.94) with p set to 0.7, and 1500 variable features identified by *FindVariableFeatures* were used as model training features. All individual daughter cells of HSC1 regardless of whether they were paired or not, were subsequently classified as HSC1, EMPP6, EMPP5, MPP Ly-I, or MPP1-4 by the classifier. Symmetric self-renewal was defined as a pair of PDCs in which both cells were classified as HSC1, whereas asymmetric self-renewal was defined as a pair in which one PDC was classified as HSC1 and the other was not. Symmetric differentiation (both committed) was defined as a pair of PDCs in which both cells were classified as downstream progenitors.

##### Processing of published data

Published 10x genomic data of LSK cells^36^ was downloaded and processed using CellRanger (v 7.2.0). Low-quality cells were filtered out as indicated in the paper, and quality-controlled data was then normalized, scaled, and integrated with HSC1/HSC2 data as described above. Clustering was performed using *FindClusters* with Leiden algorithm. To identify each cluster, enrichment scoring of gene sets generated by bulk RNA-seq^36^ was performed as described in the “Enrichment scoring” section.

#### Bioinformatic analysis of mass spectrometry data

##### Raw data preprocessing

Raw mass spectrometry files were processed using Spectronaut software (version 17.2.230208.55965, Biognosys, Switzerland) in directDIA mode with standard parameter settings. Protein identification and quantification were performed against the mouse SwissProt reference proteome (April 2024 release), comprising 17,217 curated protein entries. Protease specificity was defined as Trypsin/P, allowing up to two missed cleavage sites. Carbamidomethylation of cysteine residues was specified as a fixed modification, while methionine oxidation and protein N-terminal acetylation were included as variable modifications. Statistical confidence was ensured by controlling the false discovery rate (FDR) at 1% across peptide-spectrum match (PSM), peptide, and protein identification levels. Quantitative analysis was based on fragment ion (MS2) signal intensities, and inter-sample normalization was performed using the MaxLFQ algorithm. Peptide and protein coverage of mass spectrometry data were summarized in **Supplemental Table S7**. Data were then subjected to NAguideR^72^ to remove samples with an NA ratio over 0.5 and proteins with NA ratio over 0.5. NA value was then imputed with seqKNN.

##### GSEA analysis

Imputed data was first log_2_-transformed and then normalized using median normalization. Batch effect was estimated using R package sva^73^ (v 3.46.0) on normalized data and regressed out in log_2-_transformed fold change calculation using limma^74^ (v 3.54.2). Proteins were next ranked based on log_2-_transformed fold change, and corresponding genes were fed into *GSEA* from clusterProfiler.

#### Tracing hematopoiesis at steady state using *Flt3*^CreER^/*R26*^tdTomato^ mice

*Flt3*-creER mice were generously provided by Dr. Bo Zhou and crossed to Rosa26 ^loxP-STOP-loxP-tdTomato^ (R26^tdT^) to generate *Flt3*^CreER^/*R26*^tdT^ mice. The Flt3-creER mice were used at the age of 8-12 weeks. Both male and female mice were used in this study. Mice were randomly assigned to the different experimental groups. All mice were maintained on a C57BL/6J background. Tamoxifen was dissolved in corn oil at 10mg/ml and administrated intraperitoneally at 50 mg/kg body weight daily for five consecutive days. Two days (initial labeling) and 28 days (4-week tracing) after the final injection, the BM cells were detected for Tomato expression on distinct cell populations respectively.

### QUANTIFICATION AND STATISTICAL ANALYSIS

The data were analyzed using Microsoft Excel (Microsoft) and GraphPad Prism 8 software, except for the single-cell RNA-sequencing data, for which detailed analysis was described in the “Single-cell transcriptomic analysis section”. All data were presented as mean ± SD. All experiments were repeated as indicated; n indicates the number of independent biological repeats. No randomization or blinding was used. No statistical method was used to predetermine sample size. When two groups were compared, a two-tailed unpaired Student’s t-test was used with following specifications: 1) if qualifying normality and homogeneity of variance, Student’s t-test was used; 2) if qualifying normality but not homogeneity of variance, Welch’s t-test was used; 3) if not qualifying normality, Mann-Whitney U test was used. When multiple groups were compared, one-way ANOVA was used with following specifications:1) if qualifying normality and homogeneity of variance, one-way ANOVA with Tukey’s correction was used; 2) if qualifying normality but not homogeneity of variance, Welch’s ANOVA test with Tamhane T2 correction was used; 3) if not qualifying normality, Kruskal-Wallis test with Dunn’s correction was used. The specific statistical test used for each experiment is described in figure legends. “Not significantly different data” are not indicated.

### Supplemental information titles and legends

**Supplemental Table S1.** Immunophenotypic definition for cell populations in this study.

| Cell population | Immunophenotypic definition |
|---|---|
| **HSPC** |  |
| HSC1 | EPCR<sup>+</sup>CD150<sup>+</sup>CD48<sup>−</sup>LSK |
| HSC2 | EPCR<sup>+</sup>CD150<sup>−</sup>CD48<sup>−</sup>LSK |
| DN | EPCR<sup>−</sup>CD150<sup>−</sup>CD48<sup>−</sup>LSK |
| EMPP6 | CD34<sup>−</sup>CD135<sup>−</sup>EPCR<sup>+</sup>CD150<sup>−</sup>CD48<sup>−</sup>LSK |
| EMPP5 | CD34<sup>+</sup>CD135<sup>−</sup>EPCR<sup>+</sup>CD150<sup>−</sup>CD48<sup>−</sup>LSK |
| MPP Ly-I | CD34<sup>+</sup>CD135<sup>+</sup>EPCR<sup>+</sup>CD150<sup>−</sup>CD48<sup>−</sup>LSK |
| tradHSC | CD34<sup>−</sup>CD135<sup>−</sup>CD150<sup>+</sup>CD48<sup>−</sup>LSK |
| MPP6 | CD34<sup>−</sup>CD135<sup>−</sup>CD150<sup>−</sup>CD48<sup>−</sup>LSK |
| MPP5 | CD34<sup>+</sup>CD135<sup>−</sup>CD150<sup>−</sup>CD48<sup>−</sup>LSK |
| MPP7 | CD34<sup>+</sup>CD135<sup>+</sup>CD150<sup>−</sup>CD48<sup>−</sup>LSK |
| MPP1 | CD34<sup>+</sup>CD135<sup>−</sup>CD150<sup>+</sup>CD48<sup>−</sup>LSK |
| MPP2 | CD34<sup>+</sup>CD135<sup>−</sup>CD150<sup>+</sup>CD48<sup>+</sup>LSK |
| MPP3 | CD34<sup>+</sup>CD135<sup>−</sup>CD150<sup>−</sup>CD48<sup>+</sup>LSK |
| MPP4 | CD34<sup>+</sup>CD135<sup>+</sup>CD150<sup>−</sup>CD48<sup>+</sup>LSK |
| CLP | Lin<sup>−</sup>cKit<sup>low</sup>Sca1<sup>low</sup>CD127<sup>+</sup>CD135<sup>+</sup> |
| CMP | Lin<sup>−</sup>cKit<sup>+</sup>Sca1<sup>−</sup>CD34<sup>+</sup>CD16/32<sup>low</sup> |
| MEP | Lin<sup>−</sup>cKit<sup>+</sup>Sca1<sup>−</sup>CD34<sup>−</sup>CD16/32<sup>−</sup> |
| GMP | Lin<sup>−</sup>cKit<sup>+</sup>Sca1<sup>−</sup>CD34<sup>+</sup>CD16/32<sup>+</sup> |
| MkP | Lin<sup>−</sup>cKit<sup>+</sup>Sca1<sup>−</sup>CD150<sup>+</sup>CD41<sup>+</sup> |
| **Mature lineages** |  |
| Myeloid | Gr1<sup>+</sup>Mac-1<sup>+</sup> |
| Platelet | CD41a<sup>+</sup>Ter-119<sup>−</sup> |
| B cell | B220<sup>+</sup>Gr1<sup>−</sup>Mac-1<sup>−</sup> |
| CD4<sup>+</sup> T cell | CD4<sup>+</sup>B220<sup>−</sup>Gr1<sup>−</sup>Mac-1<sup>−</sup> |
| CD8<sup>+</sup> T cell | CD8<sup>+</sup>B220<sup>−</sup>Gr1<sup>−</sup>Mac-1<sup>−</sup> |
| **B-cell subsets** |  |
| Pro B | CD43<sup>+</sup>B220<sup>+</sup>Mac-1<sup>−</sup> |
| Pre B | IgM<sup>−</sup>CD43<sup>−</sup>B220<sup>+</sup>Mac-1<sup>−</sup> |
| Immature B | IgM<sup>+</sup>CD43<sup>−</sup>B220<sup>+</sup>Mac-1<sup>−</sup> |
| Mature B | IgM<sup>+</sup>CD43<sup>−</sup>B220<sup>high</sup>Mac-1<sup>−</sup> |
| Follicular B | CD23<sup>+</sup>CD21<sup>−</sup>CD5<sup>−</sup>CD43<sup>−</sup>B220<sup>+</sup> |
| Marginal-zone B | CD23<sup>−</sup>CD21<sup>+</sup>CD5<sup>−</sup>CD43<sup>−</sup>B220<sup>+</sup> |
| **T-cell subsets** |  |
| CD4<sup>+</sup>T<sub>naive</sub> | CD62L<sup>+</sup>CD44<sup>−</sup>CD4<sup>+</sup>NK1.1<sup>−</sup>B220<sup>−</sup> |
| CD4<sup>+</sup>T<sub>cm</sub> | CD62L<sup>+</sup>CD44<sup>+</sup>CD4<sup>+</sup>NK1.1<sup>−</sup>B220<sup>−</sup> |
| CD4<sup>+</sup>T<sub>em</sub> | CD62L<sup>−</sup>CD44<sup>+</sup>CD4<sup>+</sup>NK1.1<sup>−</sup>B220<sup>−</sup> |
| CD8<sup>+</sup>T<sub>naive</sub> | CD62L<sup>+</sup>CD44<sup>−</sup>CD8<sup>+</sup>NK1.1<sup>−</sup>B220<sup>−</sup> |
| CD8<sup>+</sup>T<sub>cm</sub> | CD62L<sup>+</sup>CD44<sup>+</sup>CD8<sup>+</sup>NK1.1<sup>−</sup>B220<sup>−</sup> |
| CD8<sup>+</sup>T<sub>em</sub> | CD62L<sup>−</sup>CD44<sup>+</sup>CD8<sup>+</sup>NK1.1<sup>−</sup>B220<sup>−</sup> |

**Supplemental Table S5.** Standards of quality control for scRNA-seq data.

| Data | Standards | Cell number Before QC | Cell number After QC |
|---|---|---|---|
| HSC1 and HSC2 | **1) nFeature_RNA<sup>a</sup>:** between 1000 ~ 4000<br>**2) nCount_RNA<sup>b</sup>:** between 10,000 ~ 200,000<br>**3) pct_counts_Mito<sup>c</sup>:** ≤5% | HSC1: 184<br>HSC2: 184 | HSC1: 159<br>HSC2: 177 |
| EMPP6, EMPP5 and MPP Ly-I | **1) nFeature_RNA:** between 1000 ~ 5000<br>**2) nCount_RNA:** between 10,000 ~ 500,000<br>**3) pct_counts_Mito:** ≤3% | EMPP6: 92<br>EMPP5: 92<br>MPP Ly-I: 92 | EMPP6: 81<br>EMPP5: 78<br>MPP Ly-I: 80 |
| HSC1-derived PDCs | **1) nFeature_RNA:** between 1500 ~ 10,000<br>**2) nCount_RNA:** between 20,000 ~ 3,000,000<br>**3) pct_counts_Mito:** ≤3% | Fresh HSC1: 43<br>PDCs: 1062 | Fresh HSC1: 42<br>PDCs: 598 (199 paired) |
| EMPP6-derived PDCs | **1) nFeature_RNA:** between 1500 ~ 8500<br>**2) nCount_RNA:** between 20,000 ~ 1,100,000<br>**3) pct_counts_Mito:** ≤2.5% | Fresh EMPP6: 48<br>PDCs: 398 | Fresh EMPP6: 40<br>PDCs: 319 (140 paired) |

a, number of detected features in a cell;
b, number of total transcripts in a cell;
c, percentage of mitochondrial mRNAs.

**Supplemental Table S7.** Peptide and protein coverage of mass spectrometry data.

| Group | Reps | Peptide | Protein |
|---|---|---|---|
| HSC1 | yH_1_1 | 12541 | 2811 |
|  | yH_2_1 | 14887 | 3270 |
|  | yH_2_2 | 14806 | 3281 |
|  | yH_2_3 | 14173 | 3170 |
| EMPP6 | yE_1_1 | 11484 | 2862 |
|  | yE_2_1 | 13348 | 3170 |
|  | yE_2_2 (removed) | 9796 | 2602 |
|  | yE_2_3 | 14604 | 3221 |

**Supplemental Table S2.** Differentially expressed genes (DEGs) between EMPP6 and HSC1 cells, as well as between either two groups of EMPP6, EMPP5 and MPP Ly-I. Related to **Fig 5 and S5**. (Excel file containing additional data too large to fit in a PDF)

**Supplemental Table S3.** Full list of GSEA results between EMPP6 and HSC1. Related to **Fig. 5**. (Excel file containing additional data too large to fit in a PDF)

**Supplemental Table S4.** High-importance features for classification of each immunophenotypic cell populations. Related to **Fig. 7 and S7**. (Excel file containing additional data too large to fit in a PDF)

**Supplemental Table S6.** Mass spectrometry data of young EMPP6 and HSC1. Related to **Fig. 5**. (Excel file containing additional data too large to fit in a PDF)

## References and Notes

1. Alhaj Hussen, K., Louis, V., and Canque, B. (2024). A new model of human lymphopoiesis across development and aging. Trends in Immunology 45, 495–510. 10.1016/j.it.2024.05.007.

2. Ross, J.B., Myers, L.M., Noh, J.J., Collins, M.M., Carmody, A.B., Messer, R.J., Dhuey, E., Hasenkrug, K.J., and Weissman, I.L. (2024). Depleting myeloid-biased haematopoietic stem cells rejuvenates aged immunity. Nature 628, 162–170. 10.1038/s41586-024-07238-x.

3. Su, T.-Y., Hauenstein, J., Somuncular, E., Dumral, Ö., Leonard, E., Gustafsson, C., Tzortzis, E., Forlani, A., Johansson, A.-S., Qian, H., et al. (2024). Aging is associated with functional and molecular changes in distinct hematopoietic stem cell subsets. Nature Communications 15, 7966. 10.1038/s41467-024-52318-1.

4. Jang, G., Contreras Castillo, S., Esteva, E., Upadhaya, S., Feng, J., Adams, N.M., Richard, E., Awatramani, R., Sawai, C.M., and Reizis, B. (2023). Stem cell decoupling underlies impaired lymphoid development during aging. Proceedings of the National Academy of Sciences 120, e2302019120. doi:10.1073/pnas.2302019120.

5. Konturek-Ciesla, A., Dhapola, P., Zhang, Q., Säwén, P., Wan, H., Karlsson, G., and Bryder, D. (2023). Temporal multimodal single-cell profiling of native hematopoiesis illuminates altered differentiation trajectories with age. Cell Reports 42. 10.1016/j.celrep.2023.112304.

6. Scherer, M., Singh, I., Braun, M.M., Szu-Tu, C., Sanchez Sanchez, P., Lindenhofer, D., Jakobsen, N.A., Körber, V., Kardorff, M., Nitsch, L., et al. (2025). Clonal tracing with somatic epimutations reveals dynamics of blood ageing. Nature 643, 478–487. 10.1038/s41586-025-09041-8.

7. Wang, Y., Zhang, W., Zhang, C., Van, H.Q.T., Seino, T., and Zhang, Y. (2025). Reducing functionally defective old HSCs alleviates aging-related phenotypes in old recipient mice. Cell Research 35, 45–58. 10.1038/s41422-024-01057-5.

8. Totani, H., Matsumura, T., Yokomori, R., Umemoto, T., Takihara, Y., Yang, C., Chua, L.H., Watanabe, A., Sanda, T., and Suda, T. (2025). Mitochondria-enriched hematopoietic stem cells exhibit elevated self-renewal capabilities, thriving within the context of aged bone marrow. Nature Aging 5, 831–847. 10.1038/s43587-025-00828-y.

9. Cheng, H., Zheng, Z., and Cheng, T. (2020). New paradigms on hematopoietic stem cell differentiation. Protein & Cell 11, 34–44. 10.1007/s13238-019-0633-0.

10. Pei, W., Feyerabend, T.B., Rössler, J., Wang, X., Postrach, D., Busch, K., Rode, I., Klapproth, K., Dietlein, N., Quedenau, C., et al. (2017). Polylox barcoding reveals haematopoietic stem cell fates realized in vivo. Nature 548, 456–460. 10.1038/nature23653.

11. Kiel, M.J., Yilmaz, Ö.H., Iwashita, T., Yilmaz, O.H., Terhorst, C., and Morrison, S.J. (2005). SLAM Family Receptors Distinguish Hematopoietic Stem and Progenitor Cells and Reveal Endothelial Niches for Stem Cells. Cell 121, 1109–1121. 10.1016/j.cell.2005.05.026.

12. Kondo, M., Weissman, I.L., and Akashi, K. (1997). Identification of Clonogenic Common Lymphoid Progenitors in Mouse Bone Marrow. Cell 91, 661–672. 10.1016/S0092-8674(00)80453-5.

13. Karsunky, H., Inlay, M.A., Serwold, T., Bhattacharya, D., and Weissman, I.L. (2008). Flk2+ common lymphoid progenitors possess equivalent differentiation potential for the B and T lineages. Blood 111, 5562–5570. 10.1182/blood-2007-11-126219.

14. Inlay, M.A., Bhattacharya, D., Sahoo, D., Serwold, T., Seita, J., Karsunky, H., Plevritis, S.K., Dill, D.L., and Weissman, I.L. (2009). Ly6d marks the earliest stage of B-cell specification and identifies the branchpoint between B-cell and T-cell development. Genes & development 23, 2376–2381.

15. Adolfsson, J., Månsson, R., Buza-Vidas, N., Hultquist, A., Liuba, K., Jensen, C.T., Bryder, D., Yang, L., Borge, O.-J., Thoren, L.A.M., et al. (2005). Identification of Flt3^+^ Lympho-Myeloid Stem Cells Lacking Erythro-Megakaryocytic Potential: A Revised Road Map for Adult Blood Lineage Commitment. Cell 121, 295–306. 10.1016/j.cell.2005.02.013.

16. Forsberg, E.C., Serwold, T., Kogan, S., Weissman, I.L., and Passegué, E. (2006). New evidence supporting megakaryocyte-erythrocyte potential of flk2/flt3+ multipotent hematopoietic progenitors. Cell 126, 415–426.

17. Boyer, Scott W., Schroeder, Aaron V., Smith-Berdan, S., and Forsberg, E.C. (2011). All Hematopoietic Cells Develop from Hematopoietic Stem Cells through Flk2/Flt3-Positive Progenitor Cells. Cell Stem Cell 9, 64–73. 10.1016/j.stem.2011.04.021.

18. Buza-Vidas, N., Woll, P., Hultquist, A., Duarte, S., Lutteropp, M., Bouriez-Jones, T., Ferry, H., Luc, S., and Jacobsen, S.E.W. (2011). FLT3 expression initiates in fully multipotent mouse hematopoietic progenitor cells. Blood 118, 1544–1548. 10.1182/blood-2010-10-316232.

19. 19. Wilson, A., Laurenti, E., Oser, G., van der Wath, R.C., Blanco-Bose, W., Jaworski, M., Offner, S., Dunant, C.F., Eshkind, L., Bockamp, E., et al. (2008). Hematopoietic Stem Cells Reversibly Switch from Dormancy to Self-Renewal during Homeostasis and Repair. Cell 135, 1118–1129. 10.1016/j.cell.2008.10.048.

20. 20. Cabezas-Wallscheid, N., Klimmeck, D., Hansson, J., Lipka, Daniel B., Reyes, A., Wang, Q., Weichenhan, D., Lier, A., von Paleske, L., Renders, S., et al. (2014). Identification of Regulatory Networks in HSCs and Their Immediate Progeny via Integrated Proteome, Transcriptome, and DNA Methylome Analysis. Cell Stem Cell 15, 507–522. 10.1016/j.stem.2014.07.005.

21. Pietras, Eric M., Reynaud, D., Kang, Y.-A., Carlin, D., Calero-Nieto, Fernando J., Leavitt, Andrew D., Stuart, Joshua M., Göttgens, B., and Passegué, E. (2015). Functionally Distinct Subsets of Lineage-Biased Multipotent Progenitors Control Blood Production in Normal and Regenerative Conditions. Cell Stem Cell 17, 35–46. 10.1016/j.stem.2015.05.003.

22. Balazs, A.B., Fabian, A.J., Esmon, C.T., and Mulligan, R.C. (2006). Endothelial protein C receptor (CD201) explicitly identifies hematopoietic stem cells in murine bone marrow. Blood 107, 2317–2321. 10.1182/blood-2005-06-2249.

23. Kent, D.G., Copley, M.R., Benz, C., Wöhrer, S., Dykstra, B.J., Ma, E., Cheyne, J., Zhao, Y., Bowie, M.B., Zhao, Y., et al. (2009). Prospective isolation and molecular characterization of hematopoietic stem cells with durable self-renewal potential. Blood 113, 6342–6350. 10.1182/blood-2008-12-192054.

24. Vazquez, S.E., Inlay, M.A., and Serwold, T. (2015). CD201 and CD27 identify hematopoietic stem and progenitor cells across multiple murine strains independently of Kit and Sca-1. Experimental Hematology 43, 578–585. 10.1016/j.exphem.2015.04.001.

25. Zhou, F., Li, X., Wang, W., Zhu, P., Zhou, J., He, W., Ding, M., Xiong, F., Zheng, X., Li, Z., et al. (2016). Tracing haematopoietic stem cell formation at single-cell resolution. Nature 533, 487–492. 10.1038/nature17997.

26. Zhang, S., Ma, Y., Wang, L., Li, X., Dong, Y., Wang, J., Cheng, T., Dong, F., and Ema, H. (2022). Expansion of Quiescent Hematopoietic Stem Cells under Stress and Nonstress Conditions in Mice. Stem Cell Reviews and Reports 18, 2388–2402. 10.1007/s12015-022-10380-6.

27. Wang, X., Dong, F., Zhang, S., Yang, W., Yu, W., Wang, Z., Zhang, S., Wang, J., Ma, S., Wu, P., et al. (2018). TGF-β1 Negatively Regulates the Number and Function of Hematopoietic Stem Cells. Stem Cell Reports 11, 274–287. 10.1016/j.stemcr.2018.05.017.

28. Xie, M., Zhang, S., Dong, F., Zhang, Q., Wang, J., Wang, C., Zhu, C., Zhang, S., Luo, B., Wu, P., and Ema, H. (2021). Granulocyte colony-stimulating factor directly acts on mouse lymphoid-biased but not myeloid-biased hematopoietic stem cells. Haematologica 106, 1647–1658. 10.3324/haematol.2019.239251.

29. Zhang, S., O’Neill, A., Xie, M., Wu, P., Wang, X., Bai, H., Dong, F., Wang, J., Zhang, Q., Suda, T., and Ema, H. (2021). Lymphoid-biased hematopoietic stem cells and myeloid-biased hematopoietic progenitor cells have radioprotection activity. Blood Science 3, 113–121. 10.1097/bs9.0000000000000089.

30. Dong, Y., Bai, H., Dong, F., Zhang, X., and Ema, H. (2021). Gene knockout in highly purified mouse hematopoietic stem cells by CRISPR/Cas9 technology. Journal of Immunological Methods 495, 113070. 10.1016/j.jim.2021.113070.

31. Liu, Z., Wang, J., Ma, Y., Xie, M., Wu, P., Zhang, S., Wang, X., Dong, F., Cheng, H., Zhu, P., et al. (2024). Early megakaryocyte lineage-committed progenitors in adult mouse bone marrow. Blood Science 6, e00187. 10.1097/bs9.0000000000000187.

32. Müller-Sieburg, C.E., Cho, R.H., Thoman, M., Adkins, B., and Sieburg, H.B. (2002). Deterministic regulation of hematopoietic stem cell self-renewal and differentiation. Blood 100, 1302–1309.

33. Wilson, Nicola K., Kent, David G., Buettner, F., Shehata, M., Macaulay, Iain C., Calero-Nieto, Fernando J., Sánchez Castillo, M., Oedekoven, Caroline A., Diamanti, E., Schulte, R., et al. (2015). Combined Single-Cell Functional and Gene Expression Analysis Resolves Heterogeneity within Stem Cell Populations. Cell Stem Cell 16, 712–724. 10.1016/j.stem.2015.04.004.

34. Rodriguez-Fraticelli, A.E., Weinreb, C., Wang, S.-W., Migueles, R.P., Jankovic, M., Usart, M., Klein, A.M., Lowell, S., and Camargo, F.D. (2020). Single-cell lineage tracing unveils a role for TCF15 in haematopoiesis. Nature 583, 585–589. 10.1038/s41586-020-2503-6.

35. Pei, W., Shang, F., Wang, X., Fanti, A.-K., Greco, A., Busch, K., Klapproth, K., Zhang, Q., Quedenau, C., Sauer, S., et al. (2020). Resolving Fates and Single-Cell Transcriptomes of Hematopoietic Stem Cell Clones by *PolyloxExpress* Barcoding. Cell Stem Cell 27, 383–395.e388. 10.1016/j.stem.2020.07.018.

36. Sommerkamp, P., Romero-Mulero, M.C., Narr, A., Ladel, L., Hustin, L., Schönberger, K., Renders, S., Altamura, S., Zeisberger, P., Jäcklein, K., et al. (2021). Mouse multipotent progenitor 5 cells are located at the interphase between hematopoietic stem and progenitor cells. Blood 137, 3218–3224. 10.1182/blood.2020007876.

37. Ema, H., Morita, Y., and Suda, T. (2014). Heterogeneity and hierarchy of hematopoietic stem cells. Experimental Hematology 42, 74–82.e72. 10.1016/j.exphem.2013.11.004.

38. Sakaue-Sawano, A., Kurokawa, H., Morimura, T., Hanyu, A., Hama, H., Osawa, H., Kashiwagi, S., Fukami, K., Miyata, T., Miyoshi, H., et al. (2008). Visualizing Spatiotemporal Dynamics of Multicellular Cell-Cycle Progression. Cell 132, 487–498. 10.1016/j.cell.2007.12.033.

39. Yo, M., Sakaue-Sawano, A., Noda, S., Miyawaki, A., and Miyoshi, H. (2015). Fucci-guided purification of hematopoietic stem cells with high repopulating activity. Biochemical and Biophysical Research Communications 457, 7–11. 10.1016/j.bbrc.2014.12.074.

40. Ema, H., Takano, H., Sudo, K., and Nakauchi, H. (2000). In Vitro Self-Renewal Division of Hematopoietic Stem Cells. Journal of Experimental Medicine 192, 1281–1288. 10.1084/jem.192.9.1281.

41. Seita, J., Ema, H., Ooehara, J., Yamazaki, S., Tadokoro, Y., Yamasaki, A., Eto, K., Takaki, S., Takatsu, K., and Nakauchi, H. (2007). Lnk negatively regulates self-renewal of hematopoietic stem cells by modifying thrombopoietin-mediated signal transduction. Proceedings of the National Academy of Sciences 104, 2349–2354. doi:10.1073/pnas.0606238104.

42. Yamamoto, R., Morita, Y., Ooehara, J., Hamanaka, S., Onodera, M., Rudolph, Karl L., Ema, H., and Nakauchi, H. (2013). Clonal Analysis Unveils Self-Renewing Lineage-Restricted Progenitors Generated Directly from Hematopoietic Stem Cells. Cell 154, 1112–1126. 10.1016/j.cell.2013.08.007.

43. Yang, L., Bryder, D., Adolfsson, J.r., Nygren, J., Ma nsson, R., Sigvardsson, M., and Jacobsen, S.E.W. (2005). Identification of Lin^−^Sca1^+^kit^+^CD34^+^Flt3^−^short-term hematopoietic stem cells capable of rapidly reconstituting and rescuing myeloablated transplant recipients. Blood 105, 2717–2723.

44. Morita, Y., Ema, H., and Nakauchi, H. (2010). Heterogeneity and hierarchy within the most primitive hematopoietic stem cell compartment. Journal of Experimental Medicine 207, 1173–1182. 10.1084/jem.20091318.

45. Somuncular, E., Hauenstein, J., Khalkar, P., Johansson, A.-S., Dumral, Ö., Frengen, N.S., Gustafsson, C., Mocci, G., Su, T.-Y., Brouwer, H., et al. (2022). CD49b identifies functionally and epigenetically distinct subsets of lineage-biased hematopoietic stem cells. Stem Cell Reports 17, 1546–1560. 10.1016/j.stemcr.2022.05.014.

46. Lin, D.S., Guckes, I.A., Ghezzi, I., Sommerkamp, P., Rodríguez-Correa, E., Frigoli, E., Cerrizuela, S., Kremer, L.P.M., Bergemann, S., Aroua, N., et al. (2026). EPCR expression marks CD150(neg) deeply dormant long-term hematopoietic stem cells. Cell Stem Cell. 10.1016/j.stem.2026.09.016.

47. Nishi, K., Sakamaki, T., Nagasaka, A., Kao, K.S., Sadaoka, K., Asano, M., Yamamoto, N., Takaori-Kondo, A., and Miyanishi, M. (2025). Alteration of long-and short-term hematopoietic stem cell ratio causes myeloid-biased hematopoiesis. ELife 13, RP95880.

48. Myers, J., Worthington, A.K., Poscablo, D.M., Medina, P., Chattopadhyaya, S., Mogharrab, R., Rommel, M.G.E., and Forsberg, E.C. (2026). Aging of hematopoietic stem cells is inconsequential to progenitor cell function. Stem cell reports. 10.1016/j.stemcr.2026.102965.

49. Yokomizo, T., Ideue, T., Morino-Koga, S., Tham, C.Y., Sato, T., Takeda, N., Kubota, Y., Kurokawa, M., Komatsu, N., Ogawa, M., et al. (2022). Independent origins of fetal liver haematopoietic stem and progenitor cells. Nature 609, 779–784. 10.1038/s41586-022-05203-0.

50. Stonehouse, O.J., Biben, C., Weber, T.S., Garnham, A., Fennell, K.A., Farley, A., Terreaux, A.F., Alexander, W.S., Dawson, M.A., Naik, S.H., and Taoudi, S. (2024). Clonal analysis of fetal hematopoietic stem/progenitor cells reveals how post-transplantation capabilities are distributed. Stem Cell Reports 19, 1189–1204. 10.1016/j.stemcr.2024.07.003.

51. Kapadia, C.D., Williams, N., Dawson, K.J., Watson, C., Yousefzadeh, M.J., Le, D., Nyamondo, K., Kodavali, S., Cagan, A., Waldvogel, S., et al. (2025). Clonal dynamics and somatic evolution of haematopoiesis in mouse. Nature 641, 681–689. 10.1038/s41586-025-08625-8.

52. 52. Patel, S.H., Christodoulou, C., Weinreb, C., Yu, Q., da Rocha, E.L., Pepe-Mooney, B.J., Bowling, S., Li, L., Osorio, F.G., Daley, G.Q., and Camargo, F.D. (2022). Lifelong multilineage contribution by embryonic-born blood progenitors. Nature 606, 747–753. 10.1038/s41586-022-04804-z.

53. Shang, F., Nizharadze, T., Thiele, R., Cirovic, B., Frank, L., Busch, K., Pei, W., Feyerabend, T.B., Höfer, T., Wang, X., and Rodewald, H.-R. (2025). Multipotent progenitors with distinct origins, clonal lineage fates, transcriptomes, and surface markers yield two hematopoietic trees. Proceedings of the National Academy of Sciences 122, e2505510122. doi:10.1073/pnas.2505510122.

54. Yu, V.W.C., Yusuf, R.Z., Oki, T., Wu, J., Saez, B., Wang, X., Cook, C., Baryawno, N., Ziller, M.J., Lee, E., et al. (2016). Epigenetic Memory Underlies Cell-Autonomous Heterogeneous Behavior of Hematopoietic Stem Cells. Cell 167, 1310–1322.e1317. 10.1016/j.cell.2016.10.045.

55. Meng, Y., Carrelha, J., Drissen, R., Ren, X., Zhang, B., Gambardella, A., Valletta, S., Thongjuea, S., Jacobsen, S.E., and Nerlov, C. (2023). Epigenetic programming defines haematopoietic stem cell fate restriction. Nature Cell Biology 25, 812–822. 10.1038/s41556-023-01137-5.

56. Flohr Svendsen, A., Yang, D., Kim, K., Lazare, S., Skinder, N., Zwart, E., Mura-Meszaros, A., Ausema, A., von Eyss, B., de Haan, G., and Bystrykh, L. (2021). A comprehensive transcriptome signature of murine hematopoietic stem cell aging. Blood 138, 439–451. 10.1182/blood.2020009729.

57. Dong, F., Hao, S., Zhang, S., Zhu, C., Cheng, H., Yang, Z., Hamey, F.K., Wang, X., Gao, A., Wang, F., et al. (2020). Differentiation of transplanted haematopoietic stem cells tracked by single-cell transcriptomic analysis. Nature Cell Biology 22, 630–639. 10.1038/s41556-020-0512-1.

58. Bolger, A., and Giorgi, F. (2014). Trimmomatic: A flexible read trimming tool for Illumina NGS data. Bioinformatics 30, 2114–2120.

59. Kim, D., Paggi, J.M., Park, C., Bennett, C., and Salzberg, S.L. (2019). Graph-based genome alignment and genotyping with HISAT2 and HISAT-genotype. Nature Biotechnology 37, 907–915.

60. Putri, G.H., Anders, S., Pyl, P.T., Pimanda, J.E., and Zanini, F. (2022). Analysing high-throughput sequencing data in Python with HTSeq 2.0. Bioinformatics 38, 2943–2945.

61. Stuart, T., Butler, A., Hoffman, P., Hafemeister, C., Papalexi, E., Mauck, W.M., Hao, Y., Stoeckius, M., Smibert, P., and Satija, R. (2019). Comprehensive integration of single-cell data. Cell 177, 1888–1902. e1821.

62. Wu, T., Hu, E., Xu, S., Chen, M., Guo, P., Dai, Z., Feng, T., Zhou, L., Tang, W., Zhan, L., et al. (2021). clusterProfiler 4.0: A universal enrichment tool for interpreting omics data. The Innovation 2, 100141. 10.1016/j.xinn.2021.100141.

63. Cao, J., Spielmann, M., Qiu, X., Huang, X., Ibrahim, D.M., Hill, A.J., Zhang, F., Mundlos, S., Christiansen, L., Steemers, F.J., et al. (2019). The single-cell transcriptional landscape of mammalian organogenesis. Nature 566, 496–502. 10.1038/s41586-019-0969-x.

64. Wolf, F.A., Angerer, P., and Theis, F.J. (2018). SCANPY: large-scale single-cell gene expression data analysis. Genome Biology 19, 1–5.

65. Ushey, K., Allaire, J.J., and Tang, Y. (2023). reticulate: Interface to Python.

66. Sievert, C. (2020). Interactive Web-Based Data Visualization with R, plotly, and shiny (Chapman and Hall/CRC).

67. Kuhn, M. (2008). Building predictive models in R using the caret package. Journal of Statistical Software 28, 1–26.

68. Liaw, A., and Wiener, M. (2002). Classification and regression by randomForest. R news 2, 18–22.

69. Takano, H., Ema, H., Sudo, K., and Nakauchi, H. (2004). Asymmetric division and lineage commitment at the level of hematopoietic stem cells: inference from differentiation in daughter cell and granddaughter cell pairs. Journal of Experimental Medicine 199, 295–302.

70. Zhang, S., Morita, M., Wang, Z., Ooehara, J., Zhang, S., Xie, M., Bai, H., Yu, W., Wang, X., Dong, F., et al. (2019). Interleukin-12 supports in vitro self-renewal of long-term hematopoietic stem cells. Blood Science 1, 92–101. 10.1097/bs9.0000000000000002.

71. Wu, T., Hu, E., Xu, S., Chen, M., Guo, P., Dai, Z., Feng, T., Zhou, L., Tang, W., and Zhan, L. (2021). clusterProfiler 4.0: A universal enrichment tool for interpreting omics data. The Innovation 2.

72. Wang, S., Li, W., Hu, L., Cheng, J., Yang, H., and Liu, Y. (2020). NAguideR: performing and prioritizing missing value imputations for consistent bottom-up proteomic analyses. Nucleic Acids Research 48, e83–e83.

73. Leek, J.T., Johnson, W.E., Parker, H.S., Jaffe, A.E., and Storey, J.D. (2012). The sva package for removing batch effects and other unwanted variation in high-throughput experiments. Bioinformatics 28, 882–883.

74. Ritchie, M.E., Phipson, B., Wu, D., Hu, Y., Law, C.W., Shi, W., and Smyth, G.K. (2015). limma powers differential expression analyses for RNA-sequencing and microarray studies. Nucleic Acids Research 43, e47–e47.

