## Supplementary material for "The self-renewing lymphoid-biased hematopoietic stem cells support long-term lymphopoiesis in mice": All supplemental figures

### Figure S1

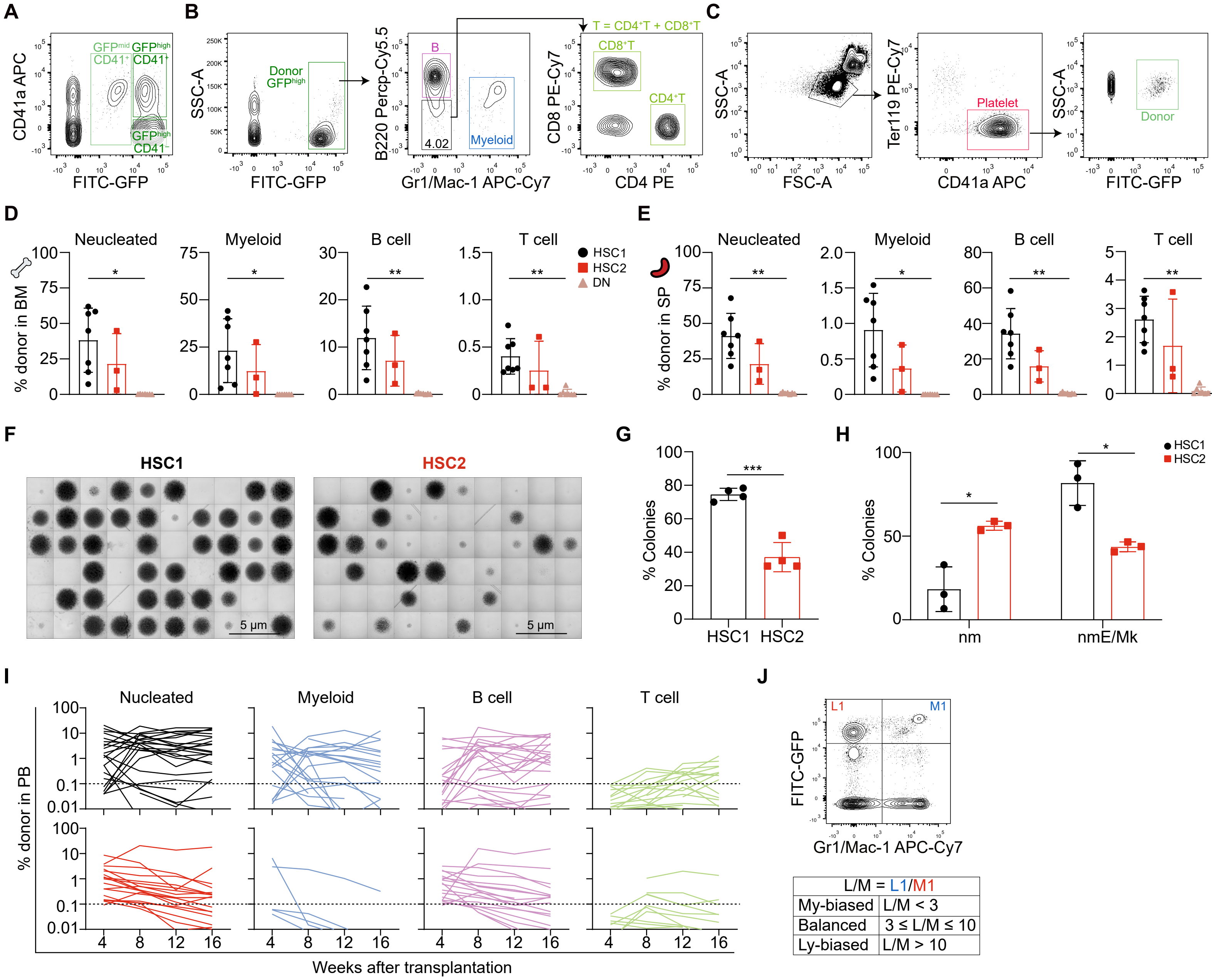
A
B
C
D
E
F
G
H
I
J
CD41a APC
FITC-GFP
GFPmid CD41+
GFPhigh CD41+
GFPhigh CD41−
SSC-A
Donor GFPhigh
B220 Percp-Cy5.5
Gr1/Mac-1 APC-Cy7
B
4.02
Myeloid
CD8 PE-Cy7
CD4 PE
T = CD4+T + CD8+T
CD8+T
CD4+T
FSC-A
Ter119 PE-Cy7
CD41a APC
Platelet
Donor
% donor in BM
Neucleated
Myeloid
B cell
T cell
HSC1
HSC2
DN
% donor in SP
HSC1
HSC2
5 μm
% Colonies
nm
nmE/Mk
% donor in PB
Nucleated
Weeks after transplantation
FITC-GFP
L1
M1
L/M = L1/M1
My-biased L/M < 3
Balanced 3 ≤ L/M ≤ 10
Ly-biased L/M > 10

### Figure S2

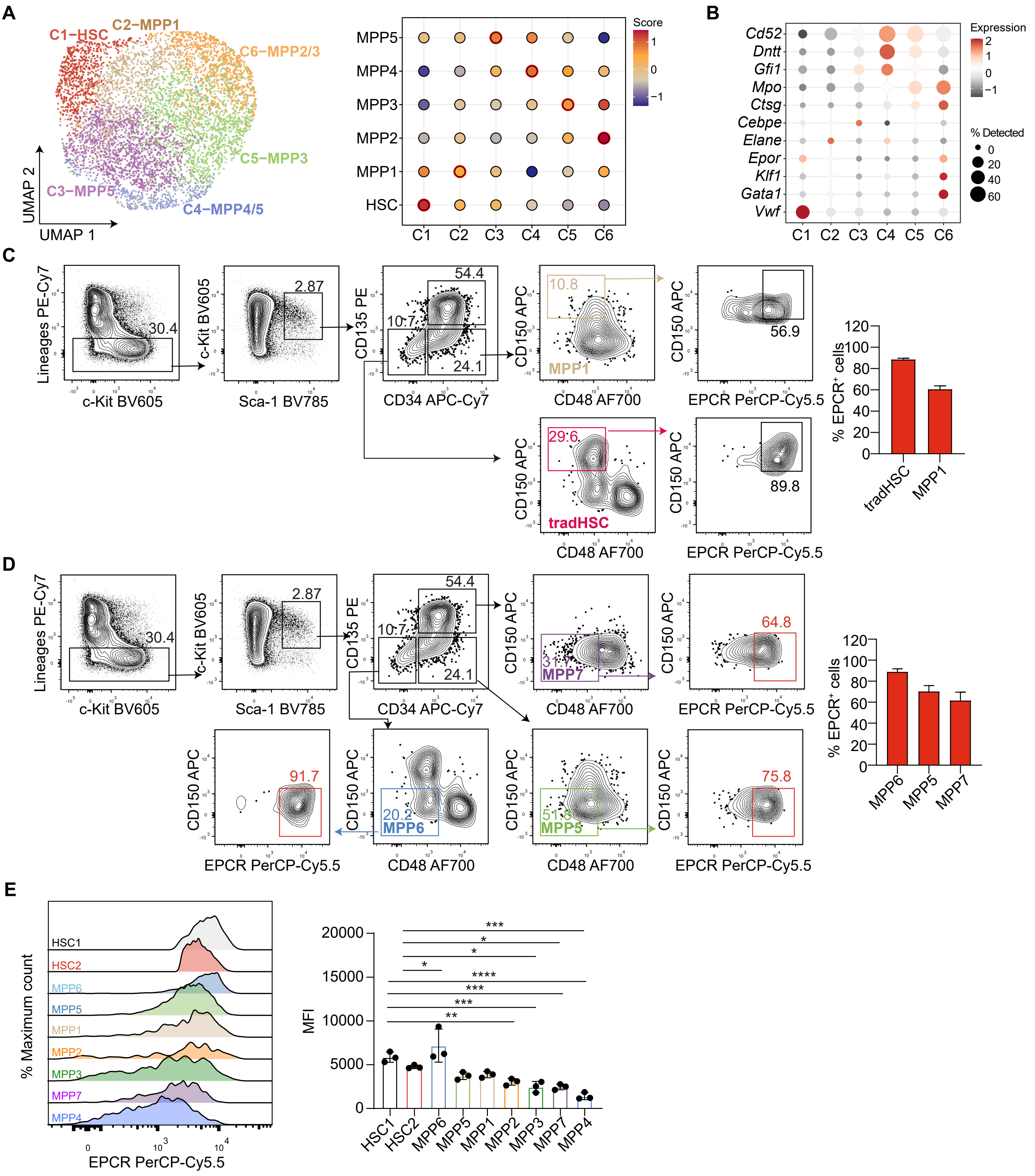
A
C1−HSC
C2−MPP1
C6−MPP2/3
C5−MPP3
C3−MPP5
C4−MPP4/5
UMAP 2
UMAP 1
MPP5
MPP4
MPP3
MPP2
MPP1
HSC
C1 C2 C3 C4 C5 C6
Score
B
Cd52
Dntt
Gfi1
Mpo
Ctsg
Cebpe
Elane
Epor
Klf1
Gata1
Vwf
Expression
% Detected
C
Lineages PE-Cy7
c-Kit BV605
Sca-1 BV785
CD135 PE
CD34 APC-Cy7
CD150 APC
CD48 AF700
EPCR PerCP-Cy5.5
MPP1
tradHSC
% EPCR+ cells
D
MPP7
MPP6
MPP5
E
% Maximum count
HSC1
HSC2
MPP6
MPP5
MPP1
MPP2
MPP3
MPP7
MPP4
MFI

### Figure S3

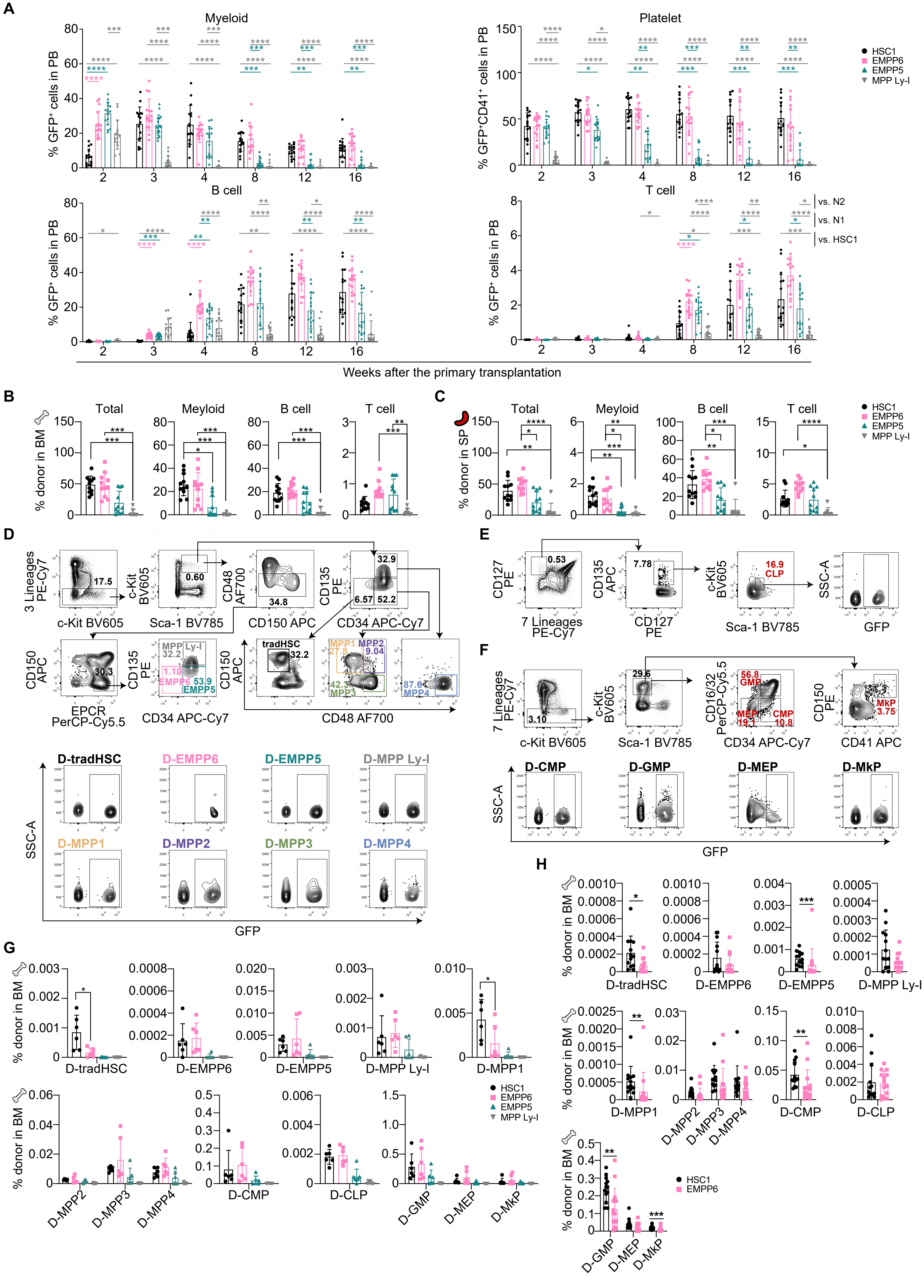

A
Myeloid
Platelet
B cell
T cell
% GFP⁺ cells in PB
% GFP⁺CD41⁺ cells in PB
HSC1
EMPP6
EMPP5
MPP Ly-I
vs. N2
vs. N1
vs. HSC1
Weeks after the primary transplantation
B
C
Total
Meyloid
B cell
T cell
% donor in BM
% donor in SP
D
E
F
c-Kit BV605
Sca-1 BV785
CD150 APC
CD34 APC-Cy7
CD48 AF700
EPCR PerCP-Cy5.5
CD127 PE
CD135 APC
CD41 APC
CD16/32 PerCP-Cy5.5
7 Lineages PE-Cy7
3 Lineages PE-Cy7
tradHSC
MPP Ly-I
EMPP6
EMPP5
MPP1
MPP2
MPP3
MPP4
CLP
GMP
MEP
CMP
MkP
D-tradHSC
D-EMPP6
D-EMPP5
D-MPP Ly-I
D-MPP1
D-MPP2
D-MPP3
D-MPP4
D-CMP
D-GMP
D-MEP
D-MkP
D-CLP
SSC-A
GFP
G
H

### Figure S4

A

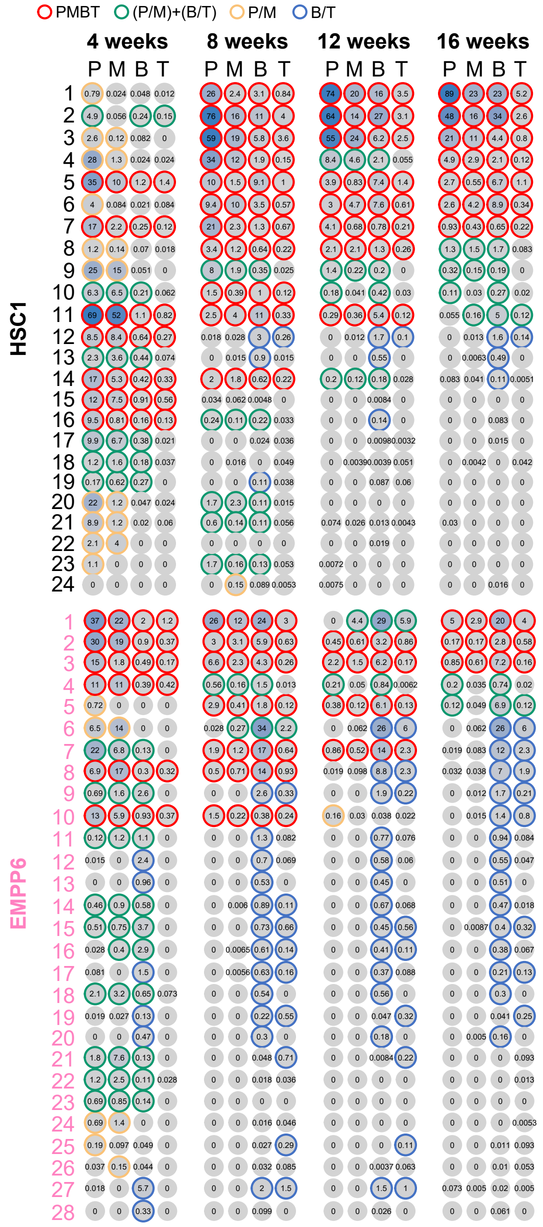
PMBT
(P/M)+(B/T)
P/M
B/T
4 weeks
8 weeks
12 weeks
16 weeks
P M B T
HSC1
EMPP6

B

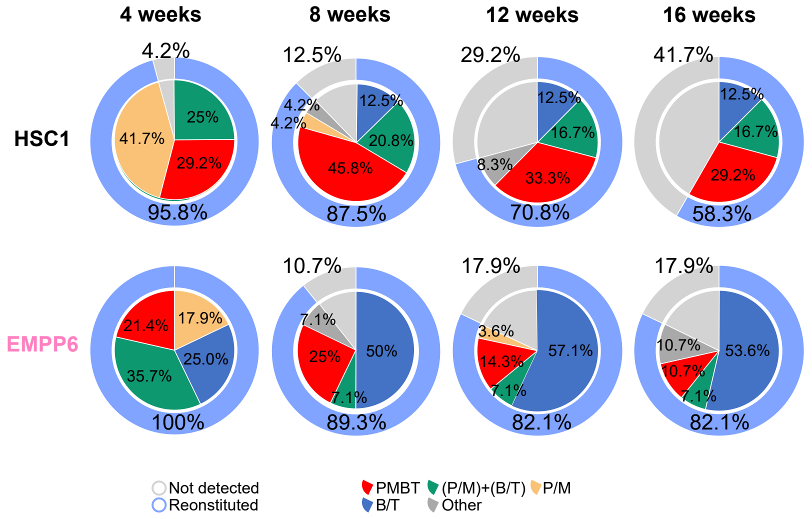
4 weeks
8 weeks
12 weeks
16 weeks
HSC1
EMPP6
4.2%
25%
41.7%
29.2%
95.8%
12.5%
4.2%
4.2%
12.5%
20.8%
45.8%
87.5%
29.2%
12.5%
16.7%
8.3%
33.3%
70.8%
41.7%
12.5%
16.7%
29.2%
58.3%
21.4%
17.9%
35.7%
25.0%
100%
10.7%
7.1%
25%
50%
7.1%
89.3%
17.9%
3.6%
14.3%
57.1%
7.1%
82.1%
17.9%
10.7%
10.7%
53.6%
7.1%
82.1%
Not detected
Reconstituted
PMBT
(P/M)+(B/T)
P/M
B/T
Other

### Figure S5

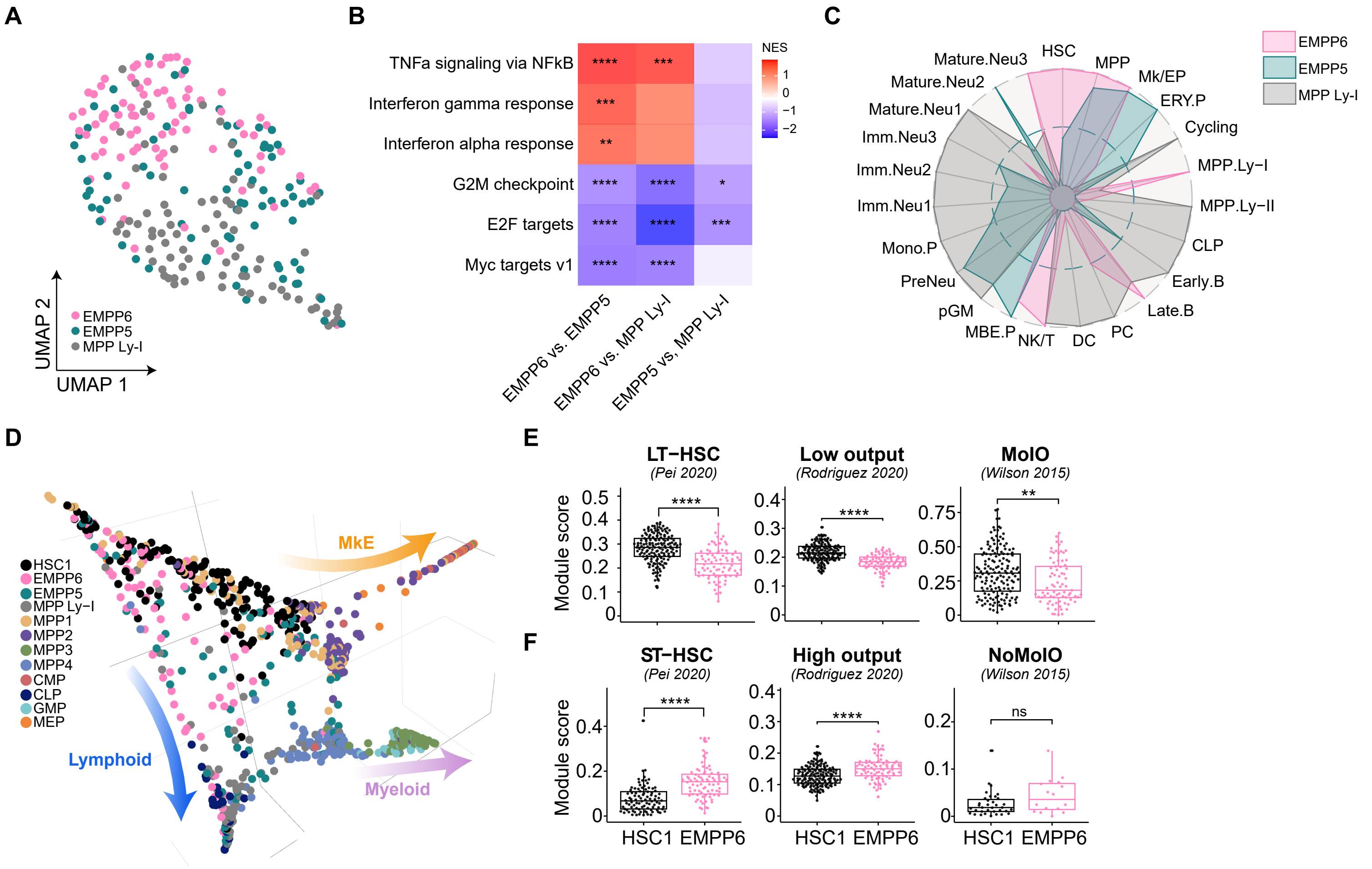
A
B
C
D
E
F
UMAP 1
UMAP 2
EMPP6
EMPP5
MPP Ly-I
TNFa signaling via NFkB
Interferon gamma response
Interferon alpha response
G2M checkpoint
E2F targets
Myc targets v1
EMPP6 vs. EMPP5
EMPP6 vs. MPP Ly-I
EMPP5 vs, MPP Ly-I
NES
HSC
MPP
Mk/EP
ERY.P
Cycling
MPP.Ly−I
MPP.Ly−II
CLP
Early.B
Late.B
PC
DC
NK/T
MBE.P
pGM
PreNeu
Mono.P
Imm.Neu1
Imm.Neu2
Imm.Neu3
Mature.Neu1
Mature.Neu2
Mature.Neu3
HSC1
MPP1
MPP2
MPP3
MPP4
CMP
CLP
GMP
MEP
MkE
Lymphoid
Myeloid
LT−HSC
(Pei 2020)
Low output
(Rodriguez 2020)
MolO
(Wilson 2015)
Module score
ST−HSC
(Pei 2020)
High output
(Rodriguez 2020)
NoMolO
(Wilson 2015)
ns
HSC1
EMPP6

### Figure S6

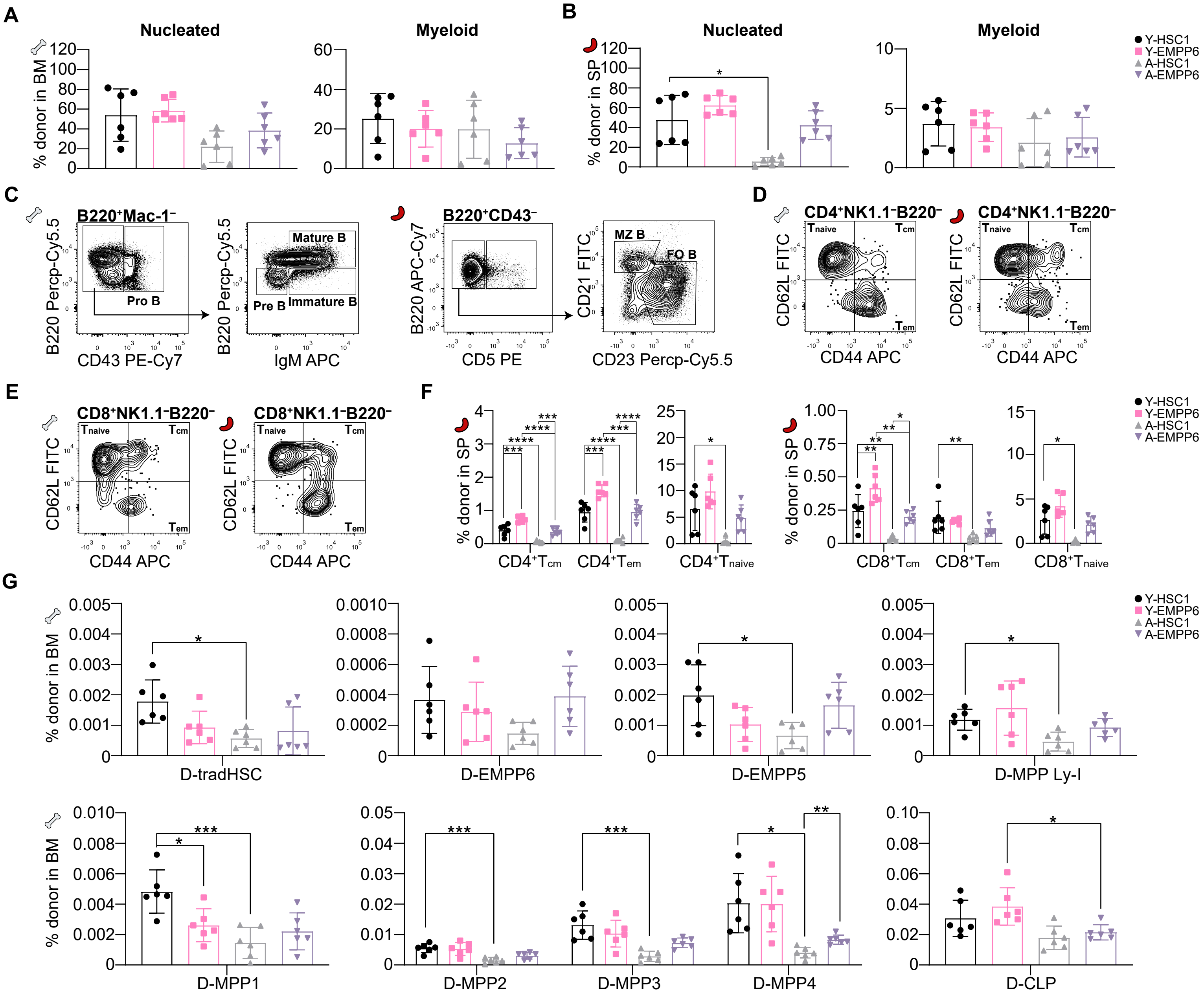

A
Nucleated
Myeloid
% donor in BM
B
Nucleated
Myeloid
% donor in SP
Y-HSC1
Y-EMPP6
A-HSC1
A-EMPP6
C
B220⁺Mac-1⁻
B220 Percp-Cy5.5
Pro B
CD43 PE-Cy7
Mature B
Pre B
Immature B
IgM APC
B220⁺CD43⁻
B220 APC-Cy7
CD5 PE
MZ B
FO B
CD21 FITC
CD23 Percp-Cy5.5
D
CD4⁺NK1.1⁻B220⁻
CD62L FITC
CD44 APC
E
CD8⁺NK1.1⁻B220⁻
F
CD4⁺Tcm
CD4⁺Tem
CD4⁺Tnaive
CD8⁺Tcm
CD8⁺Tem
CD8⁺Tnaive
Y-HSC1
Y-EMPP6
A-HSC1
A-EMPP6
G
D-tradHSC
D-EMPP6
D-EMPP5
D-MPP Ly-I
D-MPP1
D-MPP2
D-MPP3
D-MPP4
D-CLP

### Figure S7

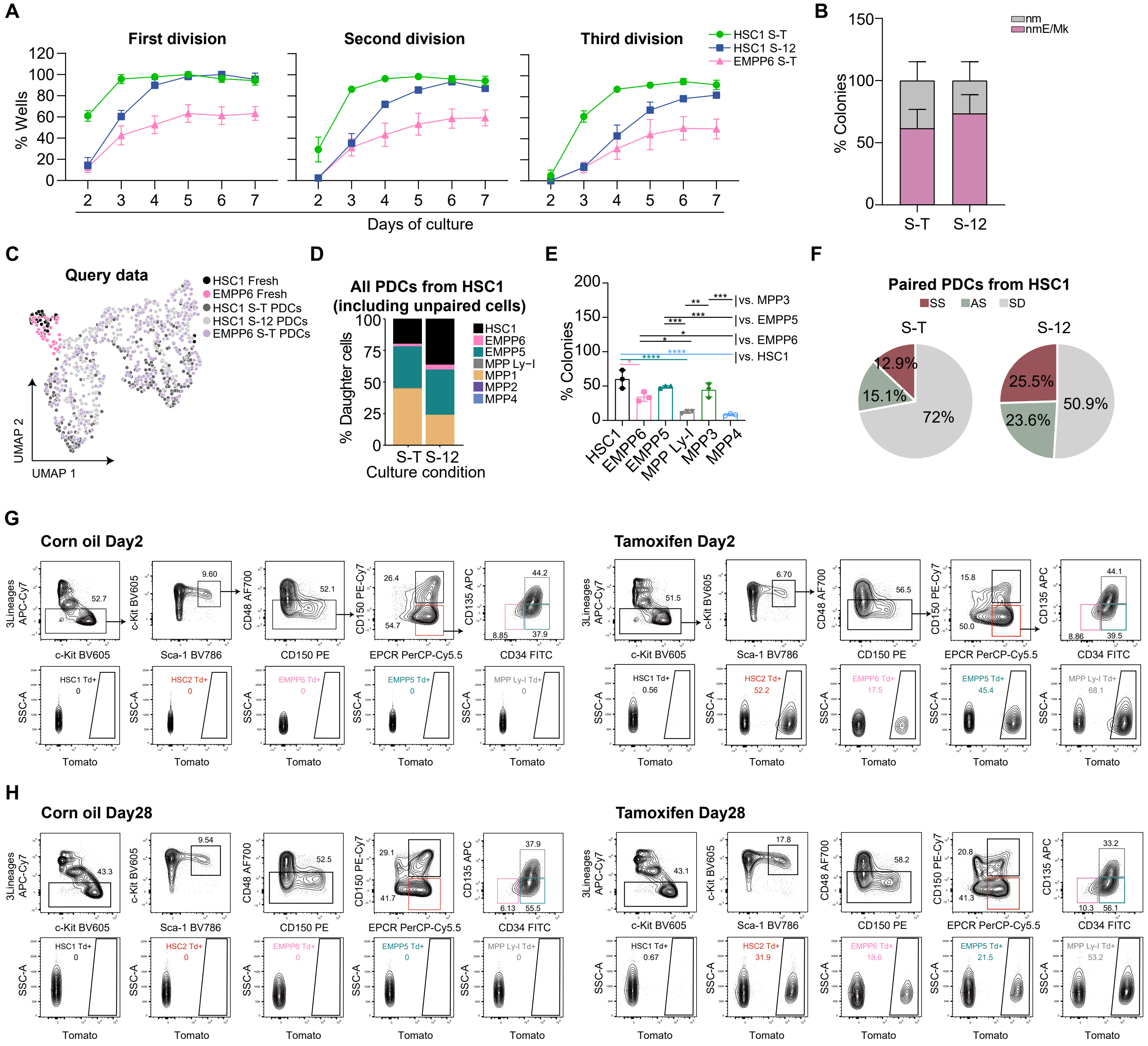

A
First division
Second division
Third division
HSC1 S-T
HSC1 S-12
EMPP6 S-T
% Wells
Days of culture
B
nm
nmE/Mk
% Colonies
S-T
S-12
C
Query data
HSC1 Fresh
EMPP6 Fresh
HSC1 S-T PDCs
HSC1 S-12 PDCs
EMPP6 S-T PDCs
UMAP 1
UMAP 2
D
All PDCs from HSC1
(including unpaired cells)
% Daughter cells
HSC1
EMPP6
EMPP5
MPP Ly-I
MPP1
MPP2
MPP4
S-T S-12
Culture condition
E
% Colonies
vs. MPP3
vs. EMPP5
vs. EMPP6
vs. HSC1
HSC1
EMPP6
EMPP5
MPP Ly-I
MPP3
MPP4
F
Paired PDCs from HSC1
SS
AS
SD
S-T
S-12
12.9%
15.1%
72%
25.5%
23.6%
50.9%
G
Corn oil Day2
Tamoxifen Day2
H
Corn oil Day28
Tamoxifen Day28
3Lineages APC-Cy7
c-Kit BV605
Sca-1 BV786
CD48 AF700
CD150 PE
CD150 PE-Cy7
EPCR PerCP-Cy5.5
CD135 APC
CD34 FITC
SSC-A
Tomato
HSC1 Td+
HSC2 Td+
EMPP6 Td+
EMPP5 Td+
MPP Ly-I Td+

### Figure S8

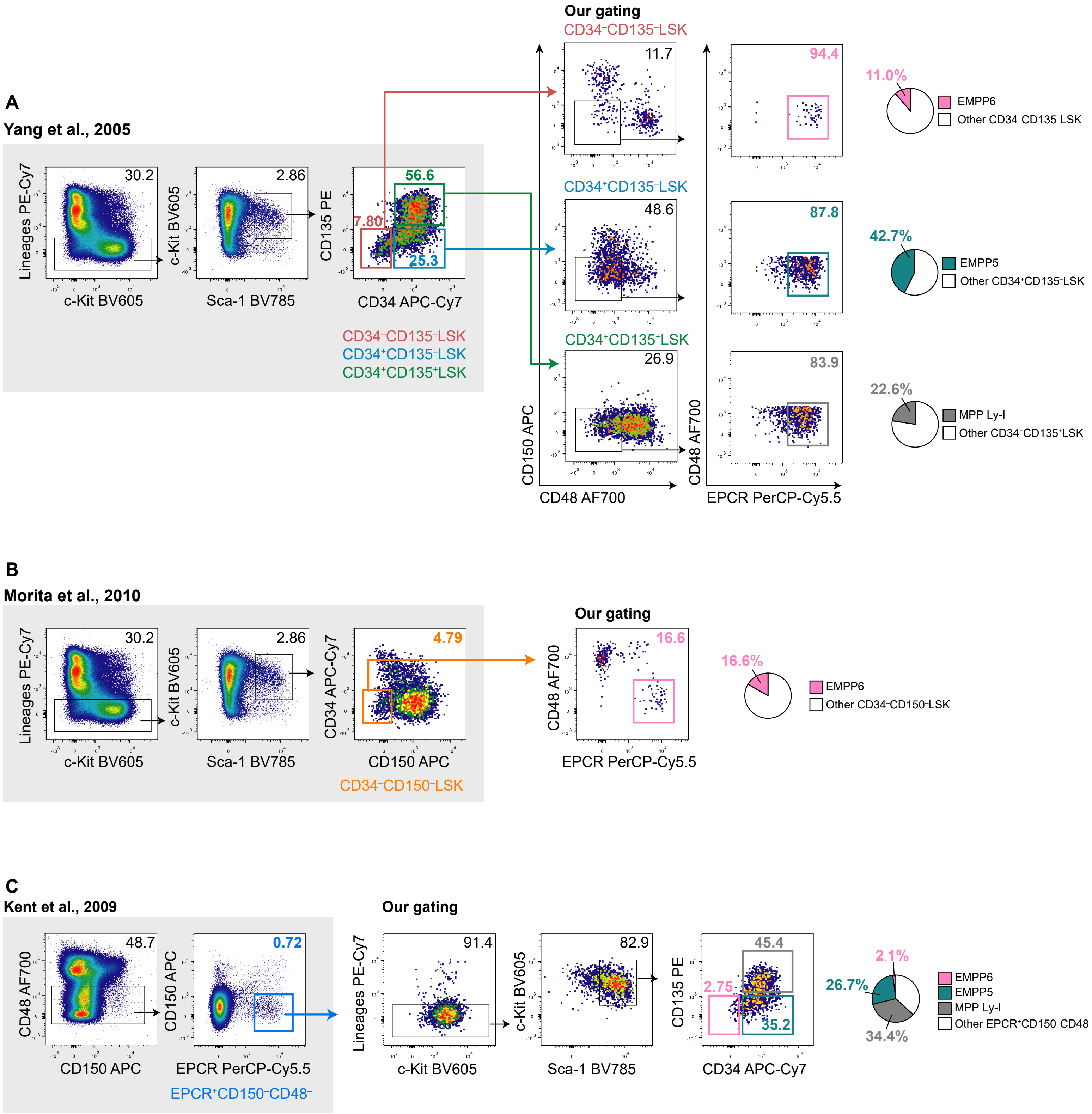
A
Yang et al., 2005
Lineages PE-Cy7
c-Kit BV605
30.2
c-Kit BV605
Sca-1 BV785
2.86
CD135 PE
CD34 APC-Cy7
56.6
7.80
25.3
CD34⁻CD135⁻LSK
CD34⁺CD135⁻LSK
CD34⁺CD135⁺LSK
Our gating
CD34⁻CD135⁻LSK
11.7
94.4
11.0%
EMPP6
Other CD34⁻CD135⁻LSK
CD34⁺CD135⁻LSK
48.6
87.8
42.7%
EMPP5
Other CD34⁺CD135⁻LSK
CD34⁺CD135⁺LSK
26.9
83.9
22.6%
MPP Ly-I
Other CD34⁺CD135⁺LSK
CD150 APC
CD48 AF700
CD48 AF700
EPCR PerCP-Cy5.5
B
Morita et al., 2010
Lineages PE-Cy7
c-Kit BV605
30.2
c-Kit BV605
Sca-1 BV785
2.86
CD34 APC-Cy7
CD150 APC
4.79
CD34⁻CD150⁻LSK
Our gating
CD48 AF700
EPCR PerCP-Cy5.5
16.6
16.6%
EMPP6
Other CD34⁻CD150⁻LSK
C
Kent et al., 2009
CD48 AF700
CD150 APC
48.7
CD150 APC
EPCR PerCP-Cy5.5
0.72
EPCR⁺CD150⁻CD48⁻
Our gating
Lineages PE-Cy7
c-Kit BV605
91.4
c-Kit BV605
Sca-1 BV785
82.9
CD135 PE
CD34 APC-Cy7
45.4
2.75
35.2
2.1%
26.7%
34.4%
EMPP6
EMPP5
MPP Ly-I
Other EPCR⁺CD150⁻CD48⁻
